# scPRINT: pre-training on 50 million cells allows robust gene network predictions

**DOI:** 10.1101/2024.07.29.605556

**Authors:** Jérémie Kalfon, Jules Samaran, Gabriel Peyré, Laura Cantini

## Abstract

A cell is governed by the interaction of myriads of macromolecules. Such a network of interaction has remained an elusive milestone in cellular biology. Building on recent advances in large foundation models and their ability to learn without supervision, we present scPRINT, a large cell model for the inference of gene networks pre-trained on more than 50M cells from the cellxgene database. Using novel pretraining methods and model architecture, scPRINT pushes large transformer models towards more interpretability and usability in uncovering the complex biology of the cell. Based on our atlas-level benchmarks, scPRINT demonstrates superior performance in gene network inference to the state of the art, as well as competitive zero-shot abilities in denoising, batch effect correction, and cell label prediction. On an atlas of benign prostatic hyperplasia, scPRINT highlights the profound connections between ion exchange, senescence, and chronic inflammation.

## Main

Understanding the cellular mechanism is considered a milestone in biology, allowing us to predict cell behavior and the impact of drugs and gene knock-outs^1–3^. A cell is regulated by a complex interplay of myriads of macromolecules that define its state. We can simplify these interactions via a gene network^4^ (GN). Many approaches have been developed to infer these networks, focusing on transcription factor (TF)-to-gene links using single-cell omics data modalities like scRNAseq and scATACseq^5–16^. This gene network subset regulating the cell gene expression levels is often called a gene regulatory network (GRN). However, many other gene products than TFs impact RNA abundances in the cell, like RNA-RNA and protein-TF interactions^17–22^. In addition, most GRN inference methods do not scale to the number of genes present in single-cell RNA datasets, and they need many cells, thus impairing their ability to reconstruct cell-state-specific networks.

Benchmarks like BeeLine^23^ and MCalla et al.^24^ have shown that despite the existence of many methods, GN inference remains a challenging problem. Indeed, it is underconstrained and has limited prior knowledge. New foundational models trained on tens of millions of measurements could help solve these difficulties. Transformers like BERT^25,26^ have gained traction in computational biology and have held promise to learn a model of the cell that would translate across many tasks of cellular biology, such as cell type annotation, batch-effect correction, perturbation prediction, and gene network inference^27^. Among them, scGPT^28^ got much attention, proposing a novel encoding of genes and their expression, a new pretraining methodology similar to autoregressive pretraining in language models, and the possibility of extracting GRN from its model (see <u>methods</u>).

Inspired by these efforts, we propose scPRINT, a foundation model designed for gene network inference. ScPRINT brings novel inductive biases and pretraining strategies better suited to GN inference while answering issues in current models (see Table S1). scPrint outputs cell type-specific genome-wide gene networks but also generates predictions on many related tasks, such as cell annotations, batch effect correction, and denoising, without fine-tuning.

We extensively benchmark scPRINT on challenging gene network inference tasks, from literature-based networks to cell type-specific ones generated via orthogonal sequencing methods. We show that scPRINT outperforms the state of the art on most of these atlas-level benchmarks. In addition, our model, focused on GN inference, is also competitive on a compendium of tasks like denoising, cell type prediction, and embedding with batch effect correction. This suggests that by learning a cell model, scPRINT gains zero-shot abilities in many tasks of cellular biology.

We use scPRINT to analyze an atlas of normal and senescent prostate tissues where we identify rare cell populations with early markers of the tumor microenvironment in B-cells. In fibroblasts, we study gene networks and recover known hubs such as PAGE4, linking the senescence of fibroblasts to changes in the ECM and downstream inflammation. We find key interconnected pathways of the oxidative stress response and extracellular matrix building via metal and ion exchange in the gene network of BPH-associated fibroblasts. We also show that healthy and disease-related cells exhibit different network patterns, demonstrating that scPRINT can help identify novel pathways and targets while considering them in their specific cellular and molecular contexts.

scPrint (https://github.com/cantinilab/scPRINT) is an open-source tool that can be readily integrated into the bioinformatics pipeline. We make public the code and model weights, but also the pretraining strategies, datasets, and our own dataloader for use with vast training sets like the cellxgene database^29^. We also release a Gene Network benchmarking suite: BenGRN and GrnnData.

## Results

### scPRINT: a scRNAseq foundation model for gene network inference

We propose scPRINT (Figure 1A), a novel bidirectional transformer designed for cell-specific gene network inference at the scale of the genome. scPRINT is trained with a novel weighted random sampling method^30^ over 40 million cells from the cellxgene^29^ database from multiple species, diseases, and ethnicities, representing around 80 billion tokens (see <u>Methods</u>). We train scPRINT at various scales (from 2M to 100M parameters) and very efficiently by using flashattention2^31^, e.g. only requiring an A40 GPU for 48 hours to train our medium model, significantly reducing the barrier to entry for any computational biology lab (see Table S2).

**Figure 1:**
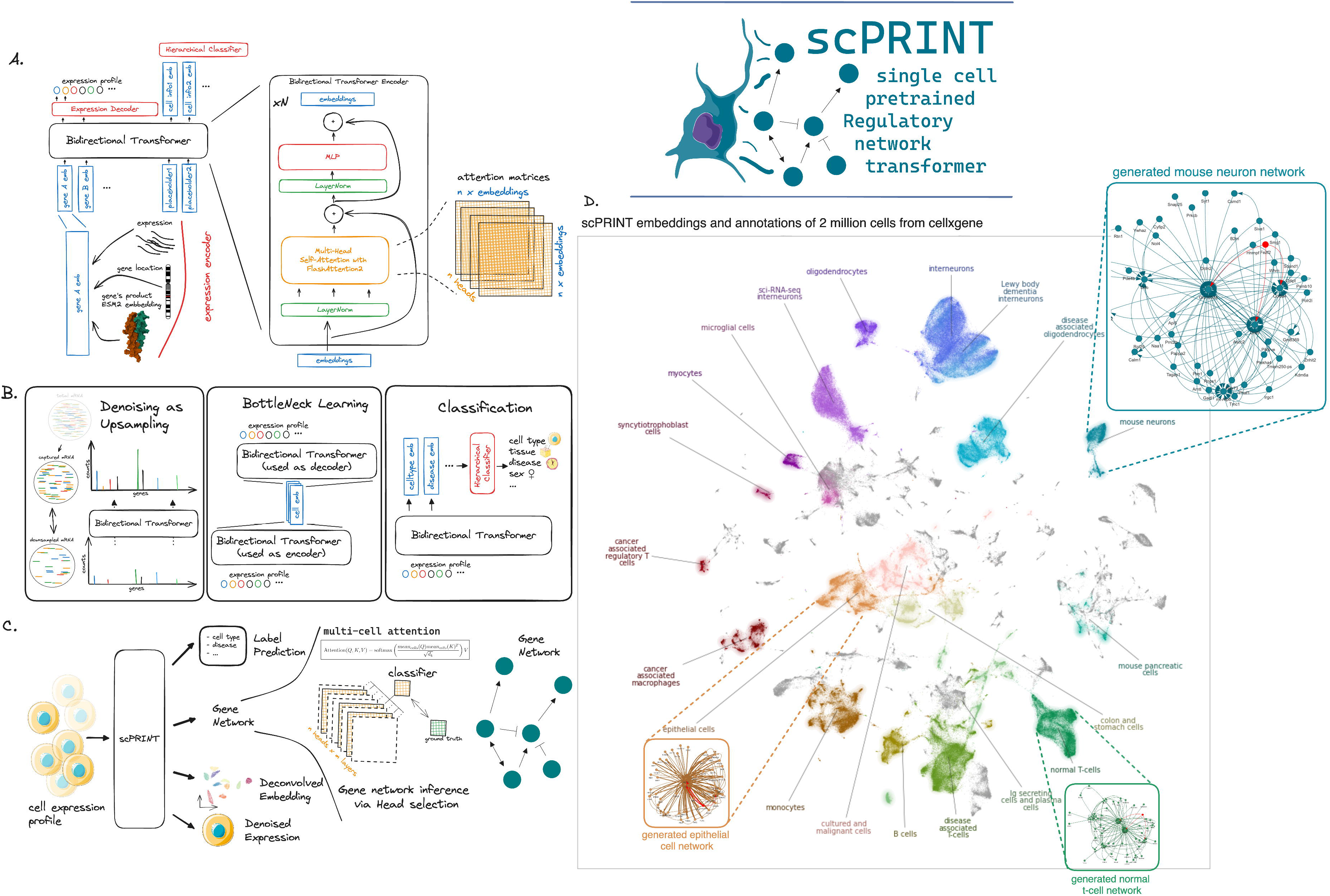
presentation of the scPRINT model and training. (a) Schematic representation of scPRINT with its bidirectional encoder, gene expression embedding encoding via gene location, matched protein ESM2 embedding, and gene expression. (b) scPRINT pre-training tasks: Denoising task whose goal is to recover the known transcriptomic profile from a purposefully downsampled expression profile. Bottleneck learning reconstructs the expression of requested genes using only their cell embedding. The same model is used for both The encoding and decoding steps. Hierarchical classification is achieved by applying a hierarchical classifier to each deconvolved embedding. This pushes the first embedding to contain cell type info, the second embedding to contain disease info, and so on (see <u>methods</u>). (c) The different outputs in scPRINT. scPRINT generates label predictions of cell type, tissue, disease, sex, sequencer, ethnicity, and organism. scPRINT generates multiple embeddings (which we call deconvolved embedding), a general one as well as a specific embedding for each class. scPRINT also generates a reconstructed expression profile at any requested sequencing depth (i.e. total transcript count) (denoising). scPRINT also generates a Gene Network by selecting and combining various attention heads into a gene x gene matrix. (d) Example of a scPRINT output from a random subset of 2 million cells from the cellxgene database. Embeddings and labels are generated by scPRINT, together with the example cell type-specific gene networks. We show only subparts of the networks extracted from a central node, represented in red.

To push scPRINT to learn meaningful gene networks (GN) and its underlying cell model, we design a novel set of pretraining tasks, as well as expression encoding and decoding schemes (Figure 1B). Similarly to ADImpute^32,33^, we expect a good gene network to help denoise an expression profile by leveraging a sparse and reliable set of known gene-gene interactions. In addition, we expect a cell model to help embed and reconstruct an expression profile by leveraging the regularities of modules and communities within its network. Finally, the cell model should represent the cell state and its different phenotypic facets. For all these reasons, we have designed a novel multi-task pre-training that combines denoising, bottleneck learning, and label prediction.

We implement the novel denoising task as the upsampling of transcript counts per cell (see <u>Methods</u>). While most other methods have been using masking as a pretraining task, our method is related to the downsampling and masking task of scFoundation^34^. We show that this strategy performs better than masked language modeling and gives scPRINT the ability to upsample any expression profile (Figure 4A).

Bottleneck learning drives scPRINT to generate a cell expression profile only from its embedding. The embedding is generated by scPRINT and is used again, this time without the cell expression profile, to regenerate the true profile (see <u>Methods</u>).

Effectively, scPRINT generates not just one such embedding per cell but multiple. For the third pre-training task, a hierarchical classifier is applied separately to each sub-cell embedding to predict labels such as cell type, disease, sex, organism, ethnicity, and sequencing platform. The embeddings are thus “deconvolved”, each representing a specific facet of the cell state. Thanks to the cellxgene database requirement for complete annotations and with our novel hierarchical classifier, we have added label prediction as part of the pretraining of scPRINT. While the assumption is that in other modalities, the scarcity and noisiness of such labels make it infeasible, we show that this approach is a net positive in our case (see Table S3). Indeed, it helps us deconvolve the various cell embeddings and performs zero-shot predictions on unseen datasets. These deconvolved embeddings are opening a future possibility to perform counterfactual generation: mixing embeddings representing different facets of cell states, e.g. fibroblast + cancer + pancreas tissue + female, to generate novel unseen expression profiles.

To encode its input expression profile, scPRINT uses gene embeddings that contain some of the prior information needed for a model to infer gene-gene^35^ and gene-DNA interactions (Figure 1A). Here, a gene in a cell expression profile is converted to an embedding by summing three representations—one of the gene itself, the other of its expression, and finally, its genomic location. The gene representation uses the ESM2^36^ amino-acid embedding of the most common protein product of that gene (see Supp Figure S1). First proposed in UCE^37^, the model learns to leverage representations that can potentially apply to unseen genes or from unseen species by using information about the protein’s structure, ontology, and evolution contained in its sequence. The gene expression is embedded via a multi-layer neural network (MLP) using log-normalized counts. Finally, to help the model understand that genes that share similar locations tend to be regulated by identical DNA regions, the gene location is embedded through positional encoding. These embeddings are concatenated with placeholder cell embeddings to form the input of the transformer model.

scPRINT trains using 2,200 randomly selected expressed genes, padded with randomly selected non-expressed genes. However, scPRINT can also make inferences on more extensive sequences of genes. We train our model using some unexpressed genes, which, combined with the denoising loss, let scPRINT discriminate the true zeros from dropouts^38^. The expression decoder of scPRINT further helps model this statistic of the data. It is a zero-inflated negative binomial graphical model inspired by previous literature in single-cell RNAseq modeling^39^. Here, the loss (also used for bottleneck learning) is thus the log-likelihood of the gene expression given the distribution parameters.

As shown in Figure 1C, at inference time, scPRINT can generate multiple outputs across any scRNA-seq-like cellular profile of various mammalian species without fine-tuning. Figure 1D shows scPRINT’s prediction at the scale of an atlas of 2M randomly sampled cells from cellxgene. From its pre-training, scPRINT performs denoising, label prediction, and cell embedding without fine-tuning. However, a critical emergent output of scPRINT is its cell-specific gene networks. Following a similar approach to ESM2, we generate cell-level gene networks via the bidirectional transformer’s input-wise weighted matrices, called attention matrices. Remarkably, we made this approach scalable enough to compute gene networks from 1 to 100,000 cells at the genome scale and with commodity hardware. These networks represent the ability of scPRINT to chart a meaningful model of cell biology. They also help make it a more explainable tool for the community, showing the network assumptions made during inference. Finally, these matrices can be further fine-tuned through classification to better reflect connections of interest (Figure 1C). For example, using literature-based gene-gene connections or perturbation-based signals.

Similarly to what has already been done in ESM2 and the Large Language Model literature^40–42^, we deeply investigate the meaning of attention matrices in the context of cellular biology, an aspect under-studied in the literature of foundation models applied to genomics.

In the following sections, we benchmark scPRINT on gene network inference against scGPT and GENIE3. scGPT is an easy-to-use, highly cited, and published transformer model^28^.

GENIE3^43^, generating a network via regression by finding the set of genes that best predict another gene’s expression, is one of the top-performing and most used methods for GRN inference.

### scPRINT recovers biological features in its gene networks

We now benchmark scPRINT against the state-of-the-art based on whether their recovered networks contain meaningful biological knowledge. We consider that a meaningful gene network should have some of its hub nodes being TFs. TFs should be more connected to their known target, on average. We should recover known gene-gene connections and expect enrichment of cell type-specific marker genes in the network.

We noticed that depending on cell type and datasets, the different tools could vary in the similarity of their GNs to Omnipath^44^. Because of this, we focused our benchmark on three randomly selected test datasets of kidney, retina, and colon tissues comprising 26 cell types^45– 47^ (see <u>Methods</u>, per dataset results in Supp. Figure S2). Of note is that we could not determine if these datasets were used during the training of scGPT.

We build one network per cell type, using 1024 cells and their 5000 most differentially expressed genes. We evaluate the quality of the networks based on their overlap with Omnipath. We also compute the network enrichment for cell type markers, TFs, and ENCODE TF targets^48^ using the prerank^49^ algorithm (Figure 2A).

**Figure 2:**
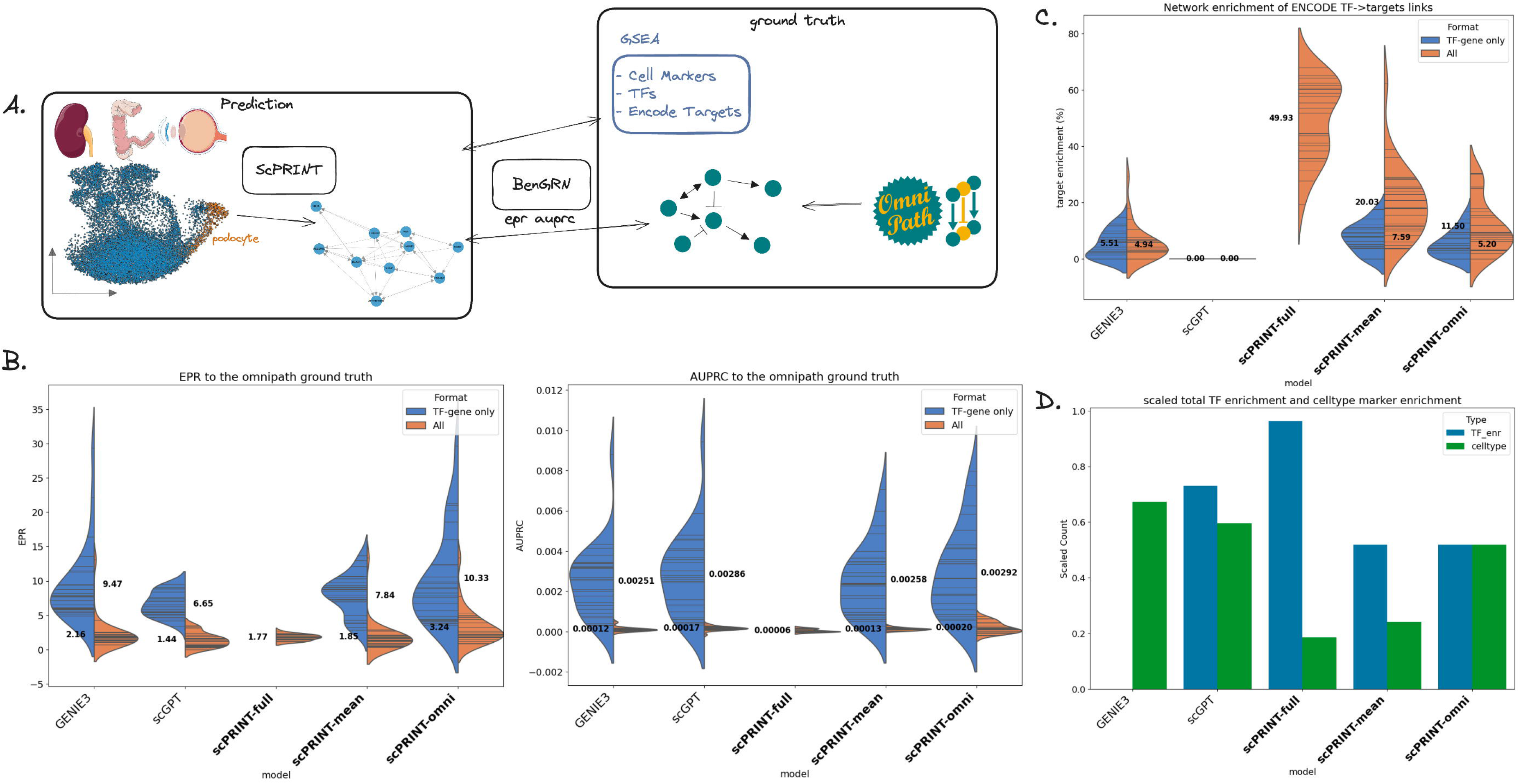
Analysis of the gene networks generated by scPRINT. (a) We extract cell type-specific gene networks for each cell type in the dataset. We perform GSEA^49^ on the network’s nodes. We compute the ability of the edges to recover the Omnipath ground truth’s connections. (b) Violin plot of the ten different AUPRC and EPR values obtained when comparing the inferred cell type-specific networks with the Omnipath network for scPRINT-full (with its genome-wide gene network), scPRINT-mean (average of all attention heads) scPRINT-omni (average of heads selected with our classifier-based method), scGPT, and GENIE3 (when considering only TF-gene connection or all gene-gene connections). (c) Violin plot of the average number of TF with enrichment for their ENCODE target in each cell type-specific network. (d) Number of GNs with a significant enrichment of TFs and of their cell type’s marker genes.

Although the scGPT code mentions GRN inference only using perturb-seq data, we reapply the same method without the perturbation-baseline comparison. This is to make it comparable with other benchmarked methods and because most of our datasets are not perturbartion-based. Similar to what is presented in its paper, we use the mean of the attention matrices across cells and the four attention heads of the last layer of the human pre-trained model. We retain this method across our benchmarks for scGPT.

For scPRINT, we generate three network versions: *scPRINT-mean*, based on the average of all heads in the model. *scPRINT-omni*, based on the average of the heads selected with our head selection method inspired by ESM2. *scPRINT-full*, which uses our method to generate genome-wide networks (see <u>Methods</u>). Indeed, in transformer models, the choice of attention heads is important. Although transformers can learn the causal structure of their input, it has been shown that some attention heads, especially in larger networks, can become unused, containing predominantly random connections^50^. Some work has been done at pruning these heads^51^ or forcing a head selection mechanism during inference and training^52^. For *scPRINT-omni*, we select heads based on a linear classifier’s prediction of the best head combination to predict Omnipath (see <u>Methods</u>). To perform this selection, we carefully split the dataset into train/test and select, using 50% of the ground truth on the first cell type of each dataset and reusing the same combination of heads across all other cell types. This shows that our selection process builds consistent networks across cell types and parts of the ground truth.

First, we look at how much information from Omnipath is contained in the inferred networks. Omnipath^44^ contains around 90,000 curated gene-gene connections, mainly from the literature. These connections are cell type agnostic, and most are TF - gene. On this benchmark, we evaluate the networks based on AUPRC and EPR, two metrics often used in GRN benchmarks^23^ (see <u>Methods</u>), where we define our task as a binary classification of connections on all gene-gene pairs. Due to the row-wise normalization of networks generated by both scPRINT, scGPT, and GENIE3, and because Omnipath has many sources with only a few targets (see Supp Figure 2), we here use the transpose of our inferred networks when making comparisons with Omnipath (see <u>Methods</u>).

In Figure 2B, we can see that *scPRINT-omni* outperforms both GENIE3 and scGPT on average across all cell types. Interestingly, all versions of scPRINT and GENIE3 outperform scGPT on the EPR metric, showing that their top predicted edges more closely match the ground truth. AUPRC results are very low overall because we do not expect most Omnipath connections to be present in the cell type’s gene network, as many connections in Omnipath might only be true in some cellular contexts. Moreover, we do not expect most connections in our generated network to exist in Omnipath as it only contains a small fraction of all real gene-gene connections. Although overall AUPRC values are small, we can see that both transformers outperform GENIE3 in the number of connections recovered. Indeed, on average, scGPT and scPRINT respectively recover 42% and 67% more connections than GENIE3.

However, GENIE3 is often used by biasing the model to generate a TF-gene graph (called GENIE3-TF, see <u>Methods</u>). This type of network, usually called a gene regulatory network (GRN), is most often used, given the importance of TFs in regulating gene expression. To compare the transformer models to GENIE3-TF, we also implement a “GRN” version of scPRINT and scGPT by subsetting their network to TF-gene connections. In this context, all the methods significantly improve their predictions without altering their relative performances. This is unsurprising, considering that Omnipath has a strong bias towards TF-gene interactions.

Interestingly, we have seen that smaller scPRINT models containing fewer heads perform better when taking the average of their heads. In contrast, head selection is often more advantageous in larger models with more heads (see Table S4). As presented at the beginning of the results section, it might be that as models become larger and less regularized, some heads tend to become unused and contain mostly noise. As a consequence, a head selection is advantageous in larger models.

We also expect biologically meaningful gene networks to have their central nodes enriched for TFs. In addition, because these networks are cell type-specific, we expect their central nodes to be enriched for some marker genes of their associated cell types (see <u>Methods</u>). In this regard, both transformer models achieve very similar and strong network enrichment for TFs compared to GENIE3, whose networks are not enriched for TFs (Figure 2C).

Moreover, amongst the 178 cell types we have marker gene sets for in pangaloDB^53^, all methods find some enrichment, especially GENIE3 and scGPT (see <u>Methods</u>). We notice that selecting heads based on Omnipath significantly improves scPRINT’s network enrichment for cell-type markers. Of note, our goal is not to annotate cell types from the gene network but mainly to showcase the cell type specificity of the networks.

Finally, we also examine how much the connections of each TF are enriched for that TF’s target. Here, scPRINT overperforms all other methods (Figure 2D). In the *scPRINT-mean* networks, 20% of the Transcription Factors for which we have data on ENCODE have connections significantly enriched for their ENCODE-validated gene targets^54^. Interestingly, only our large cell model achieved a great performance, and scGPT did not display any enrichment across the 26 cell types assessed. While we acknowledge that ENCODE is used in the Omnipath database, we cannot expect Omnipath to represent the ENCODE targets. Indeed, it combines and processes 57 additional data sources to build its consensus network.

*scPRINT-full* has been added despite its performances not being comparable to other models. Indeed, comparing its overlap with Omnipath is unfair as it includes many more genes and connections, many of which will have almost no data on this ground truth. While *scPRINT-full* showcases our ability to generate genome-wide networks, it also shows strong performances in TF enrichment and ENCODE TF-target enrichments. This highlights that even at such a large scale, networks generated by scPRINT are enriched in biological knowledge gained solely from its pre-training tasks.

Overall, we have shown that scPRINT’s cell type-specific gene networks are biologically meaningful. We will now examine cell type-specific ground truths extracted from orthogonal experiments.

### scPRINT outperforms GENIE3 and scGPT on cell type-specific ground truths

Although we have shown that our networks represent meaningful biology, the Omnipath literature-based ground truth does not reflect the topology of a biological network and is not cell type-specific. Here, we use two different modalities, perturb-seq^55^, and ChIP-seq^56^, as ground truths to compare predicted gene networks against.

In the MCalla et al.^24^ ground truth, ChIP-sequencing and perturb-seq are intersected to get at the small subset of possibly direct connections between TFs and genes for both human and mouse embryonic stem cells (ESC) (Figure 3A, see <u>Methods</u>). We have seen that these ground truth networks show a different pattern than literature-based networks (see Supp Figure S3). Some TFs regulate only a few genes, whereas others are highly connected.

**Figure 3:**
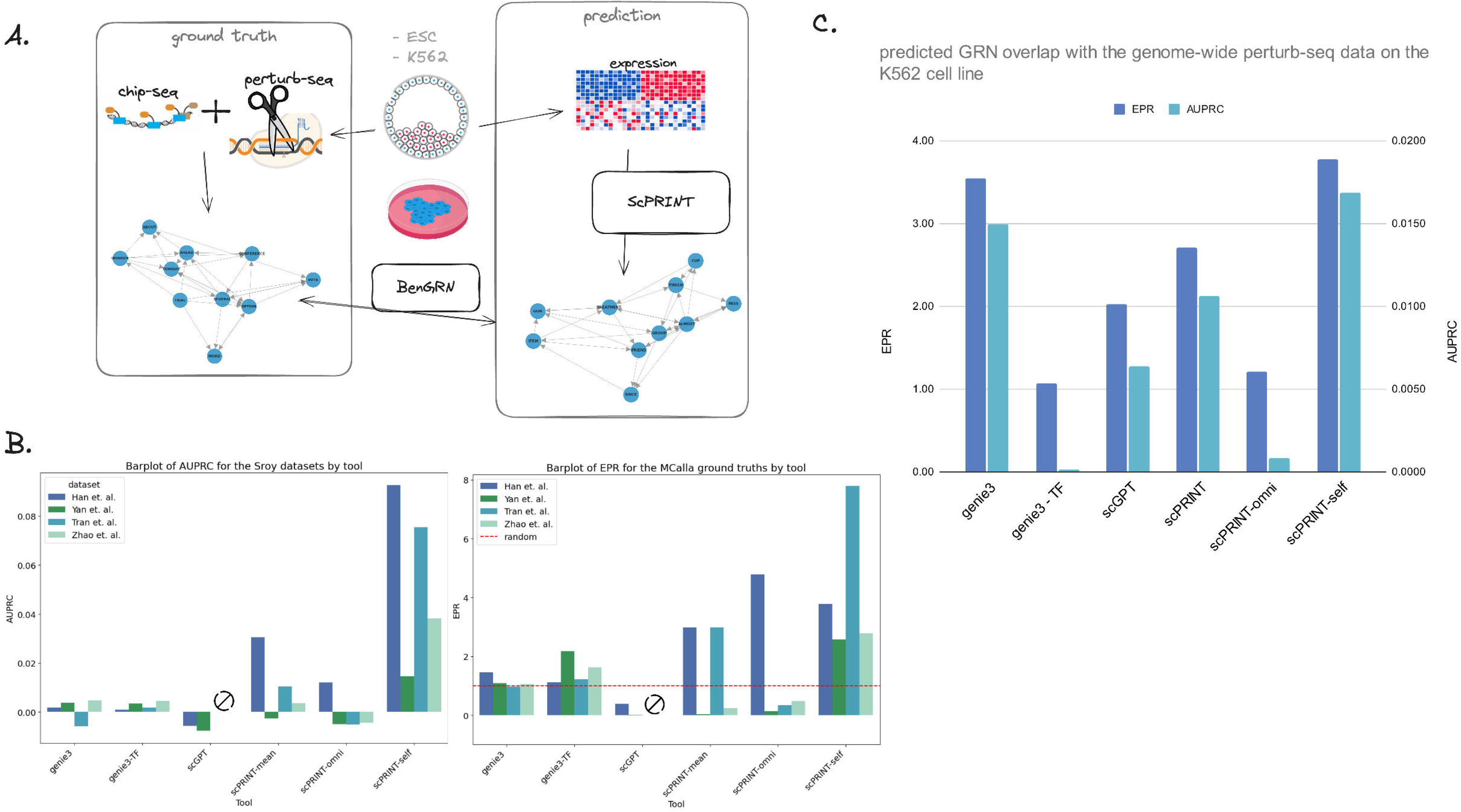
scPRINT GN inference performance on cell-type specific ground truths. (a) The ground truths are generated via orthogonal sequencing assays on the same cell type. ChIP-seq and perturb-seq are intersected for the MCalla et al. dataset on hESCs and mESCs, whereas perturb-seq on K562 is only used for the genome-wide perturb-seq ground truth. (b) Performance of scPRINT compared to GENIE3 and scGPT on the MCalla et al. ground truth using the AUPRC and EPR on two human and two mouse ESC datasets. (c) Same as (b) but on the genome-wide perturb-seq dataset. EPR and AUPRC are provided here in one barplot, left to right.

To generate our networks, we use as input one human and two mouse ESC scRNA-seq datasets from MCalla et al. with the addition of another human dataset from Yan et al.^57^. For scPRINT, three networks have been generated: one averaging all the attention heads (*scPRINT-mean*), one averaging heads selected based on how well they predicted Omnipath ground truth data (*scPRINT-omni*, for more details see scPRINT recovers biological features in its gene networks), and one averaging heads selected from the MCalla ground truth itself (*scPRINT-self*). We do not add the genome-wide network versions here as the ground truth networks are smaller, and we only assess the classification performances.

Contrary to Omnipath, some elements in these biological networks are highly connected whereas many others display no connections. This imbalance means that a model predicting only the highly connected TFs will perform well on the MCalla et al. benchmark. As a consequence we are not transposing the attention matrix as done in the previous section.

Based on both AUPRC and EPR, scPRINT outperforms GENIE3 and scGPT on this benchmark (Figure 3B). Moreover, using the TF-gene-only version of GENIE3 shows little performance gains on these ground truths, meaning that GENIE3, although only using TFs, is not selecting the right ones amongst the set of a few dozen assessed in MCalla et al.. Compared to GENIE3, which makes predictions close to random guesses, scGPT—and, in a few cases, scPRINT—can have values worse than random guessing. This means the predictions are over a specific set of TFs but not necessarily the right ones (Figure 3B)

It appeared also that selecting heads based on Omnipath, although helping slightly in one instance, is not a net benefit for this dataset. This makes sense since MCalla et al. itself does not overlap much with Omnipath (see Table S5). However, selecting heads based on the ground truth itself, only using 50% of the connections available, shows substantial improvement. These same heads also show reliable behavior when using them on the second dataset of the same species.

Overall, this shows that scPRINT can better decipher “direct” from “indirect” TF-gene connections than scGPT and GENIE3, although more tests would likely be needed. However, the results also highlight that the high imbalance (i.e., TFs being not connected or highly connected) combined with the dataset size (i.e. only a few dozen TFs assessed) and the low number of cells make the results in MCalla et al. very variable. Some of this might be true biology or explained by ChIP-seq, which can be very noisy depending on the quality of its antibodies^58^.

To answer this issue, we selected another dataset: genome-wide perturb-seq (gwps)^59^. Here, we measured the effect on transcription of knocking out all expressed genes in the K562 cell line. We transformed it into a network using a cutoff of 0.05 on the significance level of each gene’s differential expression before and after the KO of each other gene. Although this does not tell us which connections are direct or indirect, we now have a much broader set of connections over thousands of genes and better statistics to assess our gene network inference models.

Interestingly, while GENIE3 shows good results, GENIE3-TF performs poorly on this ground truth (Figure 3C). While the ground truth is not biased towards TF-gene connections, we could have expected a better overlap than random guessing from GENIE-TF.

Overall, scPRINT achieves strong performances and outperforms scGPT and GENIE3 when selecting the heads containing information on the dataset. However, again, in this dataset, selecting heads based on Omnipath does not help; the small overlap between the gwps network and the Omnipath ground truth network seems likely to be the culprit (see Table S5). These overlaps show that the three ground truth networks are very different and that a different set of heads predicts each type of ground truth.

Finally, we have seen that on both MCalla and gwps, scPRINT also predicts networks that agree with the Omnipath ground truth and are again enriched for cell type markers and TFs (see Table S6, S7).

Since GNs can be seen as approximations of a cell model, we expect that when a tool has good internal cell models, it should generate meaningful results on tasks such as denoising, cell type prediction, embedding and batch effect correction, perturbation prediction, trajectory inference, and more. We will now focus on three tasks orthogonal to GN inference to compare the ability of scPRINT to the state-of-the-art.

### scPRINT is competitive on tasks orthogonal to GN inference

To test the quality of the cell model learned by scPRINT, we now consider denoising, cell type prediction, and batch effect correction as a representative set of classic scRNAseq and cellular biology benchmarks.

Similarly to our pretraining task, we simulate lower transcript count profiles and then ask scPRINT and two other state-of-the-art methods, MAGIC^60^ and KNNsmoothing2^61^, to recreate the true expression profile. We use Spearman correlation to the original gene expression profile as our metric. In Figure 4A we show the increase in correlation after denoising the downsampled profile on 3 test set datasets, composed of ciliary body, colon, and retina tissues^46,62,63^, randomly selected from cellxgene (see <u>Methods</u>).

ScPRINT is competitive with both SOTA methods, while contrary to MAGIC and KNNSmoothing2, it operates independently over each cell in the test set. We have also seen a 10% variability in denoising ability across the different datasets used (see Table S8). This was similar across all tools and possibly related to the number of genes expressed in each dataset.

However, these test cases mostly contain very similar cell states, whereas denoising is helpful in cases with rare cell types or transitory cell states that have low cell counts by default. We show that since scPRINT does not aggregate profiles over neighboring cells, it outperforms MAGIC and KNNsmoothing2 in rare cell states subsets of the datasets (respectively: pericytes microfold cells of epithelium of small intestine and microglial cells) with around 10 to 200 cells (Figure 4A, Supp Figure S4). Computing MAGIC and KNNsmoothing2 over only this rare cell population, Gives even lower performances for MAGIC and creates an error for KNNsmoothing2 (see <u>Table S8</u>). These results suggest that a good cell model, which reliably uses learned gene-gene interactions, can help denoise an expression profile.

For cell type classification, we expect scPRINT to be able to find sets of genes that can predict a cell type across multiple batches and under the high dropout rate of single-cell RNAseq. To evaluate cell type classification, we use the multi-batch benchmark pancreas dataset of openproblems and its metrics^64,65^.

While scPRINT does not train on the test dataset itself and makes predictions over hundreds of labels, it still reaches 62% classification accuracy (Figure 4B, Supp Figure S5). Interestingly, with the macro F1 score, which considers each cell type group equally regardless of its size, scPRINT achieves similar results to the state-of-the-art^65^ methods: logistic regression and xgboost. This is probably because scPRINT is not influenced by the size of cell type groups. In addition, we have noticed that scPRINT is challenged by some specific pancreatic cell types in this dataset. Indeed, scPRINT often switches the assignment of A, B, D, and E cells. Thus, when using the coarser “endocrine” label defining these cell types, we see a strong improvement in the accuracy and macro-F1 score of scPRINT, outperforming state-of-the-art methods on the latter metric.

Here, we have shown the accuracy of scPRINT independently of cell neighborhood. However, like gene marker-based methods, scPRINT can annotate cell types in novel datasets. In this context, its predictions could be smoothed and improved using majority voting over predefined cell clusters.

Finally, scPRINT predictions are given as probability vector overall cell type labels. They can be used to display the top K labels and learn about the uncertainty of the model.

Thanks to its deconvolved embeddings, scPRINT can generate cell representations that partially remove batch effects from cell profiles. On the human pancreas and lung datasets of open problems^66^, we see that, based on the scIB metrics, scPRINT shows convincing batch effects removal ability, while not on par with the SOTA methods scGEN and scVI (Figure 4C, Supp Figure S6).

Moreover, scPRINT is one of the few methods that do not train on the test dataset and do not use already annotated batch labels. When only looking at methods that do not use batch labels as prior information, e.g. SAUCIE^67^, LIGER^68^, scPRINT is the top performer. We have also noticed that the scPRINT cell embeddings preserve biological information in a competitive way to state-of-the-art methods (Figure 4D, Supp Figure S7). This also exemplifies that a reliable cell model can perform well at deconvolving the different facets of a cell expression profile and its underlying batch effect.

Overall, we have seen that scPRINT can achieve zero-shot performances on par with many famous single-cell RNAseq tools on multiple important tasks of single-cell biology, showing that our architecture and novel pre-training tasks are a powerful new foundation for large cell models.

### scPRINT highlights the role of ion exchange and fibrosis in the ECM of Benign Prostatic Hyperplasia

To showcase the ability of scPRINT, we focus on premalignant neoplasms from an atlas of two studies of human prostate tissues^69^. The data contains both normals and pre-cancerous lesions, also called benign prostatic hyperplasia (BPH), across sequencers and age groups.

Starting from post-alignment raw counts, scPRINT generates a consistent and batch-corrected embedding of the datasets (Figure 5A, Supp Figure S8). scPRINT also annotates the cell type, sequencer, sex, ethnicity, and disease type of each cell with an accuracy of 0.71, 0.99, 0.99, 0.95, and 0.85, respectively.

**Figure 4:**
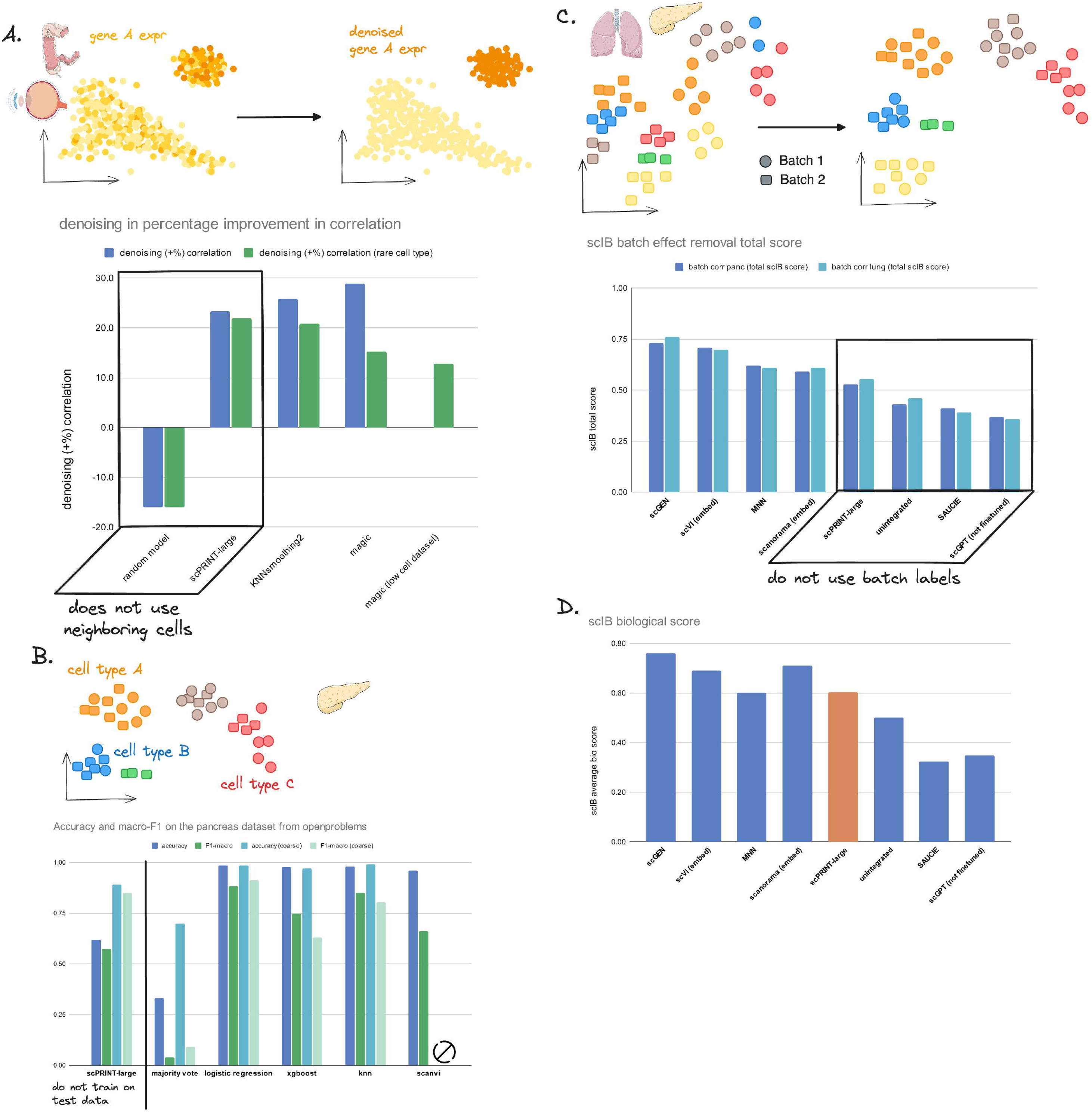
Benchmark of scPRINT on orthogonal tasks to GN inference. (a) Performance for a denoising task compared to state-of-the-art methods on a random kidney dataset from cellxgene. Here, we generate a “noisy” profile by downsampling 70% of the cell transcripts and computing the Spearman correlation increase of the correlation between the denoised and the true profile compared to the one between the noisy and the true profile. (b) Performance on cell-type label prediction compared to state-of-the-art methods. Showing accuracy, F1 and macro-F1 scores for the open-problems human pancreas dataset. (c) The performance of scPRINT compared to state-of-the-art methods on batch effect correction on the human pancreas and lung datasets from the openproblems challenge showing the scIB aggregated scores and (d) the scIB avgBIO score on both datasets

**Figure 5:**
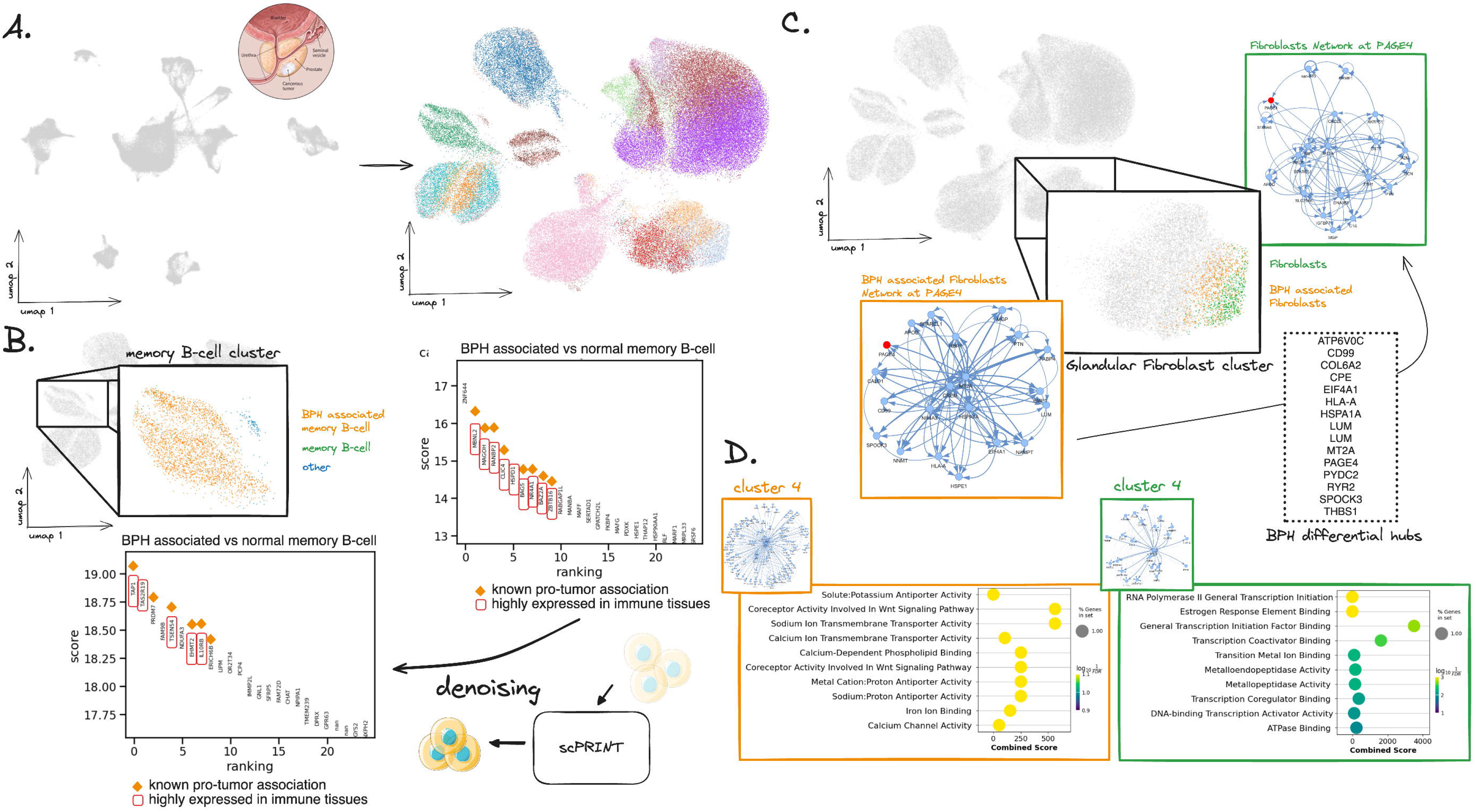
presenting a scPRINT-based bioinformatics analysis of early prostate cancer. (a) Single-cell RNAseq atlas of benign prostatic hyperplasia (BPH) and normal prostate tissues of 83,000 cells given to scPRINT. scPRINT generates a set of embeddings and label predictions for each cell. To clean our predictions, we drop cell types with less than 400 cells and diseases with less than 1000 cells, replacing them with the “other” label (see Supp Figure S8). (b) Zooming in on one cluster, we see annotations of a switched memory B-cell cluster, some labeled “benign hyperplasia” and others “normal”. Differential expression analysis on the two groups of B-cells showing enrichment of B-cell & cancer markers when assessing its top 10 genes. We performed upsampling of the transcript count before performing a new differential expression analysis where we now see new genes amongst the top 10 differentially expressed ones some of them also associated with cancer and immune tissues. (c) Zooming in on another cluster, we see annotations of a “fibroblast of connective tissue of glandular part of prostate”, some labeled as “benign prostatic hyperplasia”, and others “normal”. We generate gene networks from each and highlight a sub-network of the PAGE4 differential hub gene in BPH, showing different connection strengths and patterns between normal and BPH-associated fibroblasts. (d) Left to right: gene-set enrichment analysis, using Enrichr, of the gene community 4 found by the Louvain algorithm in the BPH-associated fibroblast gene network, same but on the normal fibroblast gene network.

We then focus on a switched memory B-cell cluster composed of a group of cells labeled as benign prostatic hyperplasia and another as normal (Figure 5A). B-cells are known to be dominant in prostate cancer and are often switched memory B-cells^70^. First, we show that they differentially express many known B-cell markers (see Supp Figure S9). In addition, when comparing the BPH to the normals B-cells, we recover that the top 10 BPH B-cells differentially expressed genes contain many known cancer markers, B cell markers, and a specific B-cell associated prostate cancer markers: *BAG5*^71^ (highlighted in Figure 5B, Table S9). Moreover, many other genes have evidence in other cancers, like *CLIC4*, known to be involved in the maintenance of the tumor microenvironment (TME) in breast cancer^72^.

However, the number of healthy cells, especially normal memory B-cells, in this dataset is small: only 26. By performing denoising, we can recover genes that might have been missed during differential expression analysis of such a low cell count. Increasing the counts of all the genes by a factor of ten and re-doing differential expression analysis highlights some new genes whose differential expression scores are even higher than those previously cited.

Interestingly amongst them, *TSEN54, EHMT2*, and *IL10RB* are known to impact the function of B-cells in malignancies (see Table S9). Other genes have evidence in immunity and cancer, like *TAP1*, which is known to be highly expressed in immune organs and is an immunomodulation gene known to play many roles in various cancers^73^, while some genes have, of yet unknown significance, like *LIP*, whose paralog *LIPA* is a known cancer target^74^ (Figure 5B).

This demonstrates how scPRINT can embed, align, and annotate diverse datasets in a meaningful way so that one can then analyze specific and rare cell clusters to recover both known and new biology.

Finally, for the second part of the analysis, we move to another cell type of interest: fibroblasts. Fibroblasts are known to be involved in cancer^75^, also called cancer-associated fibroblasts (CAFs), of which many subtypes exist, with different roles in tumor progression and invasion^76^. In our dataset, we can see a large cluster of cells labeled as “*fibroblast of connective tissue of glandular part of prostate*”, of which 500 are coming from normal tissues, and 600 are coming from hyperplasia and are possible precursors of CAFs. Interestingly here, 40% of the annotated as BPH-associated fibroblasts are coming from healthy tissue, according to the authors of the dataset. However, it is known that more than 50% of adult males over the age of 50 will have BPH^77^. Thus, one possibility is that some of the fibroblasts of these healthy tissues already present patterns of gene activation similar to those of pre-cancerous ones.

We generate a gene network of the BPH and normal fibroblasts using the 4000 most variable genes and taking the average over all heads in the network (Figure 5C). Looking at the top 15 hubs, using degree centrality, we can see S100A6 as being the top element in normal fibroblasts. This gene is known to be a fibroblast and epithelial cell marker that regulates amongst other things, cell cycle and differentiation^78,79^. We also see *MIF, IGFBP7*, and other genes involved in immune signaling and growth^80,81,82^.

However, some of these genes are not in common with the BPH fibroblasts ones. Over the set of 2881 common nodes between the two networks, the genes *HSPA1A, MT2A, SPOCK3, ATP6V0C, DEFA1, EIF4A1*, and *CD99* are considered differential hubs (i.e. more central) in the BPH fibroblasts compared to normal ones (see Table S10).

Another definition of centrality, eigenvector centrality, recovers 55% of the genes already identified as hubs, plus some new ones. As an example, Prostate Associated Gene 4 (*PAGE4*), which is part of the GAGE family of genes, is expressed in a variety of tumors and reproductive tissues, especially BPH, where it is related to oxidative stress response and fixation (i.e. anti-invasion)^83–85^. Interestingly, although the networks share 75% of their genes, they only share 50% of their edges when considering the top 20 edges per gene. It shows that over the same set of genes, scPRINT discovers distinct gene networks across biological contexts. Taking as an example the differential hub *PAGE4* (see Figure 4C), we see that it is connected to many of the top 15 hub nodes in the BPH network, such as *MT2A, HSPA1A, SPOCK3*, and *CD99*. This shows a master node sub-network linking metal and ion exchange, oxidative stress response, and inflammation^86–89^. Some genes are also part of the IL24 signaling inflammatory pathway (*EIF4A1;COL6A2;HLA-C;HSPE1*), and the secretory senescence phenotype (*H2AZ1;UBE2S;UBE2C;IGFBP7*)^80,90^, hallmarks of fibrosis and malignancies^91,92^. The *PAGE4* network in normal fibroblasts, while having some elements in common, like metal transport, is much less connected (seen by the strength of the edges in Figure 4C). It also contains a different set of genes, which are less related to senescence, inflammation, and ion exchange (see Supp Figure S10).

Furthermore, we can use these networks, defined over only a few cells, to perform community detection. Taking community 4, containing 92 genes and defined with the Louvain algorithm on the BPH-associated fibroblasts GN, we see two hub nodes: *SPOCK3* and *HERC3*. Interestingly, not much is known about those genes except that HERC3 has been linked to inflammation and the extracellular matrix (ECM) via metallopeptidase and the *NCOA1* gene^93^. SPOCK3, moreover, is known to be related to prostate malignancies and collagen in the ECM^94^. Gene set enrichment tells us that the genes in this subnetwork are mostly related to calcium, sodium, iron, and metal transport, validating the evidence around HERC3 and SPOCK3^95^. In normal fibroblast, however, taking the community most associated with metal transport (community 4, see details in Supp Figure S11 and <u>Methods</u>) shows *RNASEK, SELENOM*, and an unknown ubiquitin ligase, paralog of *ITCH*. While *RNASEK* is related to RNA degradation, its expression has been linked to a lower risk of prostate cancer^96^. *SELENOM* is of unknown function, but some SEL proteins have been related to cell adhesion^97^.

Through its networks, scPRINT highlighted the role of ion exchange and fibrosis in the ECM in Benign Prostatic Hyperplasia. While some of the same genes would have been found from differential expression analysis, these results show us how gene networks can be used to describe the intersection of genes and their molecular functions. Putting genes into the context of their connections, one can validate known functions or relate them to new ones. From such contextualization, a picture starts to emerge, whereby through specific genes, glandular fibroblasts in senescence enter a wound-healing state. This fibrosis is caused by the export of more metal and ions to generate ECM and change its acidity levels. This might cause a loss in tissue flexibility and potentially create oxidative stress^98^. In our networks, these pathways seem connected to inflammation. Chronic inflammation and wound healing states are hallmarks of BPH and a predisposition to future malignancies^99,100^

## Discussion

We can simplify the complex macromolecular interactions governing a cell through what is often referred to as a gene network. However, creating such a network in a meaningful way remains a challenging task.

We have created and benchmarked scPRINT, a novel single-cell RNA sequencing foundational model trained on more than 50 million single-cell profiles across tissues, diseases, and species contexts. scPRINT uses three novel pre-training tasks, as well as new encoding and decoding mechanisms specifically designed for gene expression data. Although it has not been directly trained for it, scPRINT generates gene networks. These networks can be used to better understand the model predictions and help make more informed decisions about the significance and role of a potential target. Finally we present a mechanism to best select heads containing the known biology of these networks. This approach also helps users fine-tune the type of network they are interested in.

We show that we outperform scGPT on many of our benchmarks while using a similar model size. We believe that our inductive biases and novel training procedures helped scPRINT achieve such a performance. Moreover, while GENIE3 is still a competitive tool, we also outperformed it on most of our benchmarks, showing that pushing training to millions of cells and large parameter sizes will be an essential direction for further work on gene network inference.

In addition, contrary to GENIE3 and scGPT, our large cell model can also achieve zero-shot performances on par with many famous single-cell RNAseq tools on multiple important tasks of cell biology. While some specialized tools might be better suited to some use cases, scPRINT’s versatility makes it a worthwhile alternative in many instances. Indeed users can directly use scPRINT in their bioinformatics workflows with commodity hardware (1 CPU, 1 GPU with 10GB of memory and 16GB of memory).

Finally, we put scPRINT to the test on a challenging atlas of normal and senescent prostate tissues showing benign prostatic hyperplasia. We identify rare cell populations with early markers of TME in B-cells. In fibroblasts, we study gene networks and recover known hubs such as PAGE4, thereby linking the senescence of fibroblasts to changes in the ECM and downstream inflammation. We find key interconnected pathways of the oxidative stress response and extracellular matrix building via metal and ion exchange in the gene network of BPH-associated fibroblasts. We also show that healthy and disease-related cells exhibit different network patterns, demonstrating that scPRINT can help identify novel pathways and targets while considering them in their specific cellular and molecular contexts.

An assumption in natural language processing is that fewer inductive biases make for better models. Our work shows that adding good inductive biases and rethinking architectures will likely be important directions for AI models in biology.

A challenging aspect of GN inference is that no perfect ground truths exist, and many GN methods are, unfortunately, benchmarked on ODE-generated mock-up expression data. In contrast, ChIP-seq, perturb-seq, and literature-based ground truths remain scarce and ambiguous. With BenGRN and GRnnData, our suite of tools for benchmarking Gene Networks inferred from single-cell RNA sequencing; we present an extensive set of real-world ground truths representative of the diversity of networks we can assess. However, improvement in performance and benchmarking will need to come from novel experimental approaches that can produce causal, genome-wide, and cell-type-specific networks.

We acknowledge that work remains to be done, from the ability of transformers to generate graphs to their explainability and the breadth of tasks they can undertake. Questions still remain regarding the pre-training tasks and how to integrate other omics modalities into foundational models.

Transcription is much more complex than what gene networks currently represent. In the future, we expect such large cell models to work in tandem with new sequencing techniques measuring modalities such as protein amounts, DNA configuration, and non-coding RNA species to solve the gap in our understanding and our ability to model cell biology.

## Methods

we propose scPRINT, a foundation model designed for gene network inference. ScPRINT brings novel inductive biases and pretraining strategies better suited to GN inference while answering issues in current models. scPrint outputs cell type-specific genome-wide gene networks but also generates predictions on many related tasks, such as cell annotations, batch effect correction, and denoising, without fine-tuning.

### Architecture

The model architecture is composed of:

- An encoder that takes the raw data and embeds it in a high-dimensional space used by the transformer.
- A bidirectional multi-head transformer
- A decoder to transform the expression embeddings into expression values
- A decoder that transforms the cell embeddings into cell-specific label prediction over a range of classes.

#### Expression encoder

In scPRINT, each gene in a cell is converted to an embedding: It corresponds to the sum of 3 different elements:

1. An embedding representing the gene itself (see Table S2 for model embedding size). ESM2^36^ embedding of each gene’s most common protein product was used to represent that gene. While imperfect in some ways, this inductive bias allows the model to learn representations that potentially apply to even unseen genes from unseen species or integrate specific genetic mutations into its representation. First implemented in UCE^37^, this provides the model information related to the gene product’s structure, ontology, and similarity to other genes. This also speeds up the training greatly, particularly for small models. We show that this is a great gene representation but that model performance can be increased by refining gene embeddings further during training. However, we elect not to do so to maintain the model’s versatility in working on unseen genes. We encode the genes’ embeddings using ESM2. The mapping process happens the following way:
  - A gene name is mapped to its canonical protein name using Ensembl^101^.
  - We recover the protein sequence of the protein using Ensembl
  - We use the protein sequence to generate an embedding using ESM2 by averaging over all the amino-acid output embeddings as done in the ESM2 paper.
  With the embedding function provided in our code, one can easily do this with any species in Ensembl. scPRINT can effectively be retrained with any set of gene embeddings, which can be frozen during training or used only for initialization (tried, for example, in our ablation studies, Table S3).
2. An embedding of the gene location in the genome. This has also been proposed in UCE and helps the model understand that genes with similar locations tend to be regulated by similar regulatory regions^102^, a relationship well-known in cellular biology. We encode the genes’ locations using positional encoding. Here, every gene less than 10,000 bp from the next is said to be in the same location; otherwise, we increment location by 1. We do this for all genes in the Ensembl database per species. We then embed these locations by applying the Positional Encoding (PE) algorithm of Vaswani et al. ^25^.
3. An embedding of the gene expression in the cell. For this, we embed the gene’s expression using an MLP. While GeneFormer came up with a ranking strategy based on a gene expression compared to a baseline expression, scGPT instead used binning of log normalized counts. On our end we haven’t found that this approach was the simplest nor was performing better than only using the log-transformed counts. We thus directly take the log-transformed counts

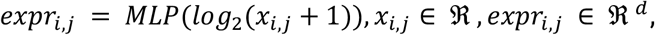

where *expr*_*i,j*_ is the embedding of the expression, *x*_*i,j*_ is the expression value of the gene j in the cell i, and the MLP is a two-layer neural network, where each layer is composed of

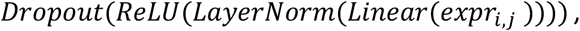

where the Dropout rate is fixed at 0.1, and the dimensions are specified as 1 → *d* for the first layer of the MLP and *d* → *d* for the second layer, with d representing the model dimension.

Finally, when encoding a cell expression profile, only a subset of 2200 genes is used during pretraining. If less than 2200 genes are expressed, we randomly choose 2200 expressed genes and pad them with randomly sampled unexpressed genes (meaning with an expression value of 0). This approach allows the model to see different patches of the same cell profile during training. We chose 2200 genes as 2/3rds of the cells in cellxgene had less than this number of genes expressed, striking a balance between computation and gene usage.

We decided to add unexpressed genes because, combined with our denoising methodology, this lets the model figure out that some genes are true 0s during training. In contrast, others are only caused by dropout and a function of the transcript counts. This causes scPRINT to model dropout as a function of read depth (i.e. total transcript count).

Moreover, this completes the minibatch by token matrix without padding and fully utilizes the GPU during the attention computation.

The full set of embeddings sent to the transformer looks the following way

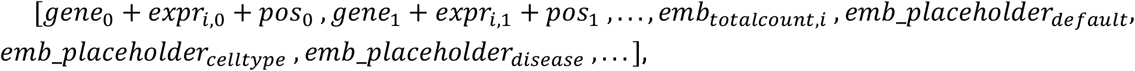

where *gene*_*j*_ is the gene j encoding, *expr*_*i,j*_ is the encoding of the expression of gene j in cell i, and *pos*_*j*_ is the gene j location encoding.

The total count information is stored separately and encoded similarly to the expression,

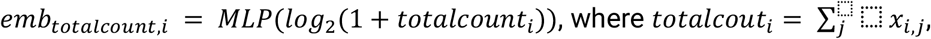

with *x*_*i,j*_ the expression value of gene j in cell i, and the MLP is a two-layer neural network similar to the previous one.

The full cell total count (totalcount) lets scPRINT model its denoising based on this required total count parameter.

The placeholder tokens (total count, default cell embedding, cell type, disease, sex, ethnicity, assay, organism) are learned embeddings that stay the same across all inputs. They only act as placeholders for the model to fill in during the forward process. At the transformer output, they will have been modified to contain the embeddings requested. At least two are used, one containing the default cell embedding and another the profile’s total depth. More tokens can be used, one for each predicted cell label.

#### Model

The model is a bidirectional autoencoder similar to BERT^26^ with *n* layers, *h* attention heads, and a dimension of *d*. It uses the flashattention2^31^ methodology implemented in Triton to compute its attention matrix. It uses the pre-normalization technique^103^, with a sped-up layer norm implemented in Triton’s tutorial^104^. It uses a stochastic depth with increasing dropout probability^105^.

It has a 2-layer MLP with a 4x width increase in its hidden layer and a gelu activation function.

#### Expression decoder

scPRINT uses a novel expression decoder for foundation models, which outputs the parameters of a zero-inflated negative binomial (*ZiNB*) function for each gene *i* in cell *j*. The *ZiNB* distribution is defined as

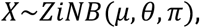

where the parameters *μ, θ, π* are obtained from a multi-layer perceptron (MLP) applied to the expression embeddings outputted by the transformer model at its last layer (*emb*_*empr*_), which are the:

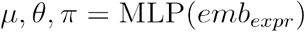

The MLP is a two-layer neural network with dimensions [*d, d*, 3]

Based on the work of Jiang et al.^38^, zero inflation is the best distribution when considering a broad range of transcriptomic measurements, where some have enough dropouts, and a zero inflation term is needed to model it. In our case, and similarly to scVI^39^, we define our *ZiNB* as

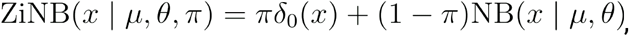

where *δ*_0_ (*x*) is a point mass at zero, and *NB*(*x* | *μ, θ*) is the negative binomial distribution with mean *μ* and dispersion *θ*.

With these parameters, the negative binomial distribution is represented in the following way

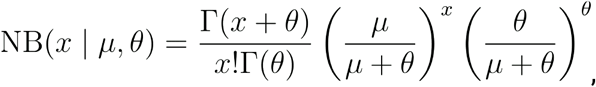

where *μ* is the mean and *θ* the overdispersion parameter, which here represents the inverse of the dispersion. From Hibe et al.^106^, we know that this is a parameter change from the most used probability mass function (PMF) given by

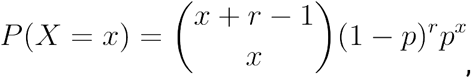

where r is the number of successes, *p* is the probability of success, and *k* is the number of failures.

One can interpret such a negative binomial distribution as a Poisson distribution with an additional overdispersion term that makes the variance not tied to the mean. In scPRINT, we use the zero-inflated Poisson for count downsampling as we can’t easily infer the gene overdispersion parameter from each cell profile. By removing this zero-inflated Poisson from the gene expression profile, we keep the potential overdispersion in the profile (see the Negative Binomial to Poisson relationship section in Methods).

Compared to scVI, where the overdispersion parameter *θ* is learned for each gene, we make scPRINT output it together with *μ, π* (see Supp. Figure S12)

Effectively, the model learns that the dispersion might change depending on the gene, the sequencer, the cell type, and the sequencing depth.

#### Class decoder

scPRINT also outputs a variety of class embeddings, such as default cell embedding, cell type embedding, disease embedding, etc., by filling the different placeholder tokens given as input (see the Expression encoder section in the Methods).

Effectively, for each class, we have the model learn to produce a new deconvolved embedding (e.g. cell type, disease, tissue, age). This means the model uses an MLP to transform each token where A is a class. For each, we jointly train a classifier

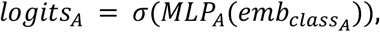

Where:

- *logits*_*A*_ represents the logits for a class A of a dimension *d*_7_ whose size corresponds to the number of labels.
- *σ* denotes the Sigmoid activation function.
- *MLP*_*A*_stands for the Multi-Layer Perceptron trained to predict the logits of the class *A*.
- 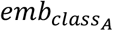 is the output embedding for the class A of dimension *d*.

However, some classes, like cell type, have up to 800 labels. Fortunately, cellxgene classes follow an ontology, a robust structure that defines relationships among the labels. We reduce the size of the output labels by training the model only on the leaf labels in the ontology hierarchy (i.e. the most precise available). For cell types, this represents around 400 different labels (see Table S11).

Thus, when a label is not very specific for a cell type (e.g. neuron), the model will predict the best leaf label (e.g. dopaminergic neuron). This way, we can generate meaningful training signals from even very coarse labels (see <u>The classification task</u> section in methods for more information and definition of the loss). We only apply this hierarchical classifier to the cell type, disease, and assay labels.

In the following section, we show how we train such classifiers. During the classifiers’ training, we sum up their loss without applying any scaling between the different classes.

### Pretraining

The three tasks of the multi-task pretraining are the denoising task, the classification task, and the bottleneck learning task. While the denoising loss enhances the model’s ability to find meaningful gene-gene connections, the other two try to make the model and its underlying networks more robust and cell-type-specific. All three losses are summed without rescaling.

#### Optimization method

The optimization is done with fused ADAMW, with a weight decay of 0.01. We noticed a total inability to learn when using base ADAM, which has a similar weight decay. This can be explained by a known inequivalence issue in ADAM^107^.

We use the stochastic weight averaging^108^ method during training with a learning rate of 0.03. During pre-training, the hyperparameters are set to dropout of 0.1, a learning rate (LR) of 1e-4, the precision is set to 16-mixed with residuals in fp32. We clip gradients to 100 and train in many sub-epochs of 7000 training batches and 2000 validation batches with a warmup duration of 500 steps.

Across epochs, we use a linear LR decrease of 0.6 with a patience of 1 and stop training after three consecutive increases in validation loss (patience: 3). In the final layer of the class decoders, we initialize values to a normal distribution around 1 for weights, 0 for biases, and - 0.12 for biases.

Our batch size is 64, and we use a pre-norm strategy for the transformer with a linearly increasing stochastic depth dropout rate of 0.02 per layer. We use a noise parameter of 60%. We split the cells in the datasets into 98% train and 2% validation and reserve at minimum 2% of separated datasets for testing.

Finally, we use weighted random sampling on our training data based on the different class values we have to predict. We use a factor of 50, meaning the rarest elements will, on average, be sampled only 50 times less than the most common ones. The sampling factor used for each group is then 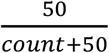, instead of 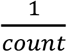 where count is the number of cells in each group.

#### The classification task

We perform label prediction during pretraining for different classes, currently: cell type, disease, sequencer, ethnicity, sex, and organism. Due to issues in the ontologies, we have omitted tissue type and age classes.

Due to the hierarchical structure of the prediction, we also created a hierarchical loss. Here, we compute the loss regularly when the label is a leaf label. Otherwise, we replace all associated leaf labels to the given label by the log-sum-exp, such that for a cell label, the loss is:

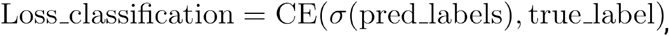

with:

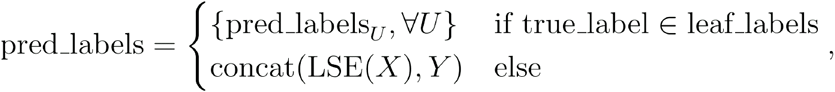

where:

- *Y* = {*pred_labels*_*U*_, ∀ *pred_labels*_*U*_ ∉ *node*_*group*}
- *X* = {*pred*_*labels*_*U*_, ∀ *pred*_*labels*_*U*_ ∈ *node*_*group*}
- True _labels being a vector of size *n*_*labels*_ one hot encoded from true_label
- pred_labels a vector of size *n*_*leaf_labels*_
- node_group is the subset of the labels that share the common parent node true_label when true_label ∉ leaf-labels.

The CE (cross-entropy) is defined as:

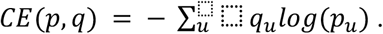

And the LSE (log-sum-exp) is defined as

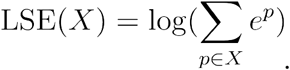

This loss allows the classifier to learn even in cases where the labels can be of varying coarseness without the coarseness of some labels impacting the ability of the model to predict the true fine-grained labels (see Supp. Figure S13)

The loss is hierarchical for the classes: cell type, disease, sequencer, ethnicity; the labels follow a hierarchy defined by Cell Ontology, MONDO, EFO, HANCESTRO^109–112^, respectively.

We do not compute the loss for cells where a class has an unknown label. We perform these classification tasks in one pass, using the embeddings generated directly from the downsampled expression profile.

#### The denoising task

Similarly to ADImpute, we expect a good gene network to help denoise an expression profile by leveraging a sparse and reliable set of known gene-gene interactions. In addition, we expect a good cell model to help embed and reconstruct an expression profile by leveraging the regularities of modules and communities within its network.

We view denoising similarly to upsampling, and inversely, we view adding noise as downsampling a cell profile.

Noise is similar to downsampling because of the distribution we are working with. Note that contrary to vision tasks (e.g. diffusion models), where additive Gaussian noise is added, in the context of expression data, where the distribution is often seen as a Poisson, NB, or ZINB, the data is already noisy, and the more counts are sampled, the less noise. No information is similar to not sampling data.

We downsample an expression profile using a zero-inflated Poisson model of the data. With this formulation, on average, half of the counts to be dropped are dropped by randomly removing a number of reads per gene, given by sampling from a Poisson whose lambda parameter is proportional to the number of counts in that gene. The remaining half of the counts to be dropped are dropped by randomly setting some genes to 0, i.e. a complete dropout of that gene. It is to be noted that with this definition of downsampling, the exact average amount of counts dropped for both parts depends slightly on the dropout *r*. During our pretraining, *r* is set to 0.6, meaning, on average, 60% of the transcript counts are dropped per cell.

Let *x*_*i*_ be the gene expression vector of cell i with dimensions *n*_*genes*_; we create a downsampled *version* by doing

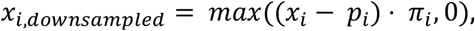

with:

- *p*_*i*_ ∼ *poi*ss*on*(*x*_*i*_ *× r ×* 0. 55) a vector of size *n*_*genes*_ where the poisson is samples for each element *x*_*i*_ of x
- *π*_*i*_ = *I*(*u* ≥ *r* × 0.55) a vector of size *n*_*genes*_, the binary mask vector indicating non-dropout genes.
- *u*_*i*_ ∼ *Uniform*(0,1), a vector of size *n*_*genes*_. of random values drawn from a uniform distribution.
- · denotes the element-wise multiplication.
- *r* being the dropout amount. We scale it by a tuning hyperparameter of 0.55 instead of 0.5 for numerical reasons.

The goal of the model is then, using *x*_*I,downsam*_ as an input, to output the parameters *μ*_*i*_, *θ*_*i*_, *π*_*i*_ of a *ZINB* distribution of the true profile *x*_*i*_, all vectors of size *n*_*genes*_. The contribution of cell i to the loss is then computed as the negative log-likelihood of the count data given the distribution parameters being generated by the model

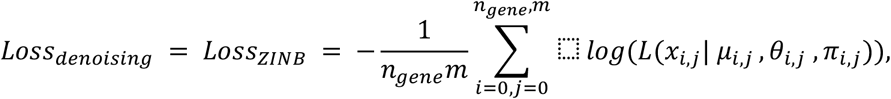

where *n*_*genes*_ is the size of the expression profile *x*_*i*_, m is the size of the minibatch and

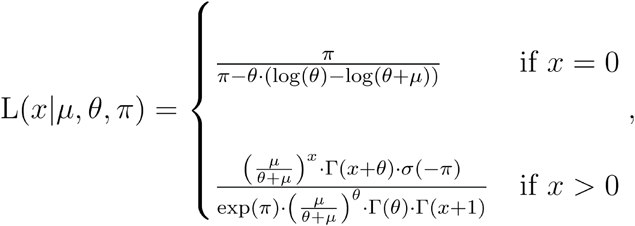

with *σ* the sigmoid function.

We show that models trained with such a framework perform better than regular MSE-trained models (see Table S3), for which one only outputs one value instead of three, which directly represents the log-transformed count of the data. In this case, the loss is the mean squared error between the predicted and true count values.

scPRINT effectively lets the user choose between the three formulations: *ZINB* with a *ZINB* loss, NB with an NB loss, and direct log-transformed count reconstruction with an *MSE* loss.

However, we have noted that the *NB* and *ZINB* loss still have some notable issues. They can easily overflow, especially when working with lower precision systems (like fp16, bf16, …). These losses are also proportional to the total expression count, meaning cells with higher expression will have a higher loss on average. It also appears that the log-likelihood cannot go below ∼1.1 loss on average and plateaus quickly. This makes evaluation of the loss less practical when comparing models. Finally, this minimal loss also depends on the total number of zeros in the true expression dataset, as the zero-inflation part of the loss converges smoothly to 0.

#### The bottleneck learning task

Bottleneck learning is a method that drives the model to generate a cell expression profile only from its embedding. Cell-embedding that can be passed again to that same model without the gene expression information, such that from the cell-embedding only, scPRINT can re-generate the cell’s expression profile. The model thus finds the best compression of the cell’s expression according to the information-theoretic theorem by Tishbi et. al.^113^.

While many transformer models and Geneformer directly use the average of gene embeddings to generate a cell embedding, this will likely squash the expression information.

scGPT used another methodology (called “MVC”) to generate an embedding vector such that

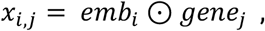

where *x*_*i,j*_ is the expression of gene j in cell i, and *⊙* is the dot product. For each gene embedding *gene*_*j*_, the embedding only contains information about the gene name, not gene expression. Regular MSE on each *x*_*i,j*_ is then used as the training loss.

This pushes the cell embedding *emb*_*i*_ to contain all the expression information of the cell i.

This is less computationally intensive to train than our bottleneck learning method. However, we have noticed poorer reconstruction through this methodology than ours (see Table S3).

In our case, we consider that our model scPRINT can act as two parts of an autoencoder. The encoding part is when we give scPRINT the expression profile of a cell and retrieve a set of deconvolved cell embeddings (see the <u>Class decoder</u> section of the methods). The decoder part is when we provide scPRINT only the gene labels without their corresponding expression values and the deconvolved cell embedding in place of the empty placeholder embeddings (see Supp Figure S14).

This means the encoder is considered as

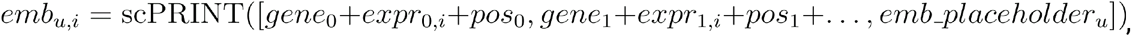

where *emb*-_*u,i*_ is the output embedding of the placeholder embedding token u for the cell i (in our case, we use multiple (default, totalcount, cell_type, disease, sex, organism, ethnicity, sequencer). Then the decoder is defined as

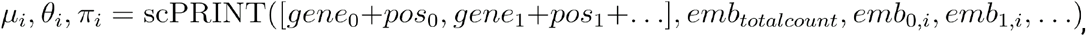

With *μ*_*i*_, *θ*_*i*_, *π*_*i*_ vectors ofsize *n*_*genes*_. Finally, the loss is given by the ZINB loss:

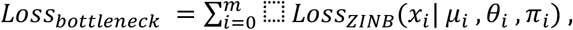

where *x*_*i*_ is the cell i expression profile and *m* the minibatch size.

Implementing a set of deconvolved embeddings is not straightforward. In our case, we push the embeddings to be as different from one another as possible with a contrastive loss defined as

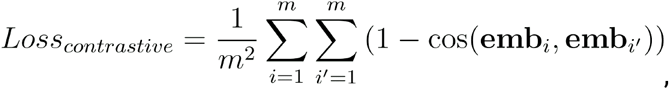

where *emb*_*i*_ and *emb*_*i*_ are the cell embeddings, *m* is the minibatch size, and *cos* denotes the cosine similarity.This pushes each embedding to represent the correct information using the classifiers. However, more is needed to remove all the batch effects or entirely prevent information leakage across embeddings.

Finally, we have also used the classifier output logits as cell embeddings. This works particularly well for cell type, disease, or sequencer classes containing many labels. It has been shown that classifier logit outputs behave similarly to embeddings^114^ and, in our case, offer an even better removal of the batch effects (see Supp Figure S6).

For the bottleneck loss, we directly reconstruct expression using the cell embeddings generated from the noisy, downsampled expression profile of the denoising process, doing the entire process in one single pass. We sum all the losses without scaling them:

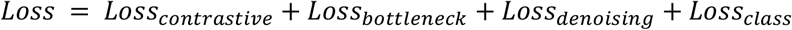

### scDataloader

Parallel to this work, we worked with Lamin.ai to develop a dataloader for large cell atlases, described and benchmarked in Rybakov et al.^30^. One key advantage of this dataloader is its ability to perform weighted random sampling on hundreds of millions of cells without being a bottleneck during pretraining. scDataloader samples cells amongst the 800+ datasets of cellxgene’s mid-2023 release, using the cell labels to inform how rare the specific combination of labels is.

From this, the dataloader produces a cell sampling weight, rescaled with a hyperparameter. The dataloader will sample, with replacement, more consistently rare cell types than more common ones.

We have produced an additional wrapper package around the laminDB “mapped-dataset” called scDataloader. scDataloader works with lamin.ai but can also interface with scVI and AnnData formats to enable downloading, preprocessing, and QC of large single-cell databases and datasets. It is very flexible and can represent expression data in the formats used by scPRINT, scGPT, and Geneformer. It also implements a lightning datamodule scheme and command line interfaces for quick setup (see Supp Figure S15).

Overall, we preprocess each of the 1200 datasets in cellxgene by only keeping primary cells from either humans or mice and dropping all the spatial omics datasets. Spatial omics are not true single-cell assays, and we decided for now not to include them. We also drop any cells with less than 200 expressed genes. Finally, we drop any resulting dataset smaller than 100 cells, with less than 10,000 genes, or from which more than 95% of the cells have been removed. This results in a new database of 54,084,961 cells and 548 datasets.

We believe that the weighted random sampling strategy allowed our pre-training to be much faster by creating more diverse minibatches.

### Extracting meta-cell gene networks from attention matrices in scPRINT

Transformers compute multiple attention matrices per layer, called attention heads. This is done by splitting the generated *K, Q*, and *V* embedding into *m* sub-embeddings, thus defining *m* attention heads. Each attention head computes the attention matrix via the equation:

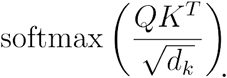

However, we would want to aggregate those over multiple cells from a similar cell state to increase the signal obtained from only one cell. We are doing so by averaging the Keys and Queries embeddings over the set of cells *U* passed to the model:

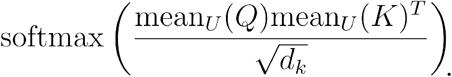

By doing this, the attention matrix behaves as if each query vector for cell i was “looking” across the key vectors of all the cells in U.

The resulting object is a row-wise normalized *n*n* matrix, where *n* is the size of the input context (i.e. the number of genes passed to the model). However, we also include the possibility to generate large matrices and gene networks, referred to as genome-wide gene networks. We take the average over different sets of expressed genes for each cell in the set U. This allows us to compute a genome-wide attention matrix while only doing forward passes on smaller subsets of the genome per cell.

### Heads selection

With scPRINT, we present a method to select heads based on some available ground truth data. This is inspired by the ESM2 paper^115^ and uses a somewhat similar method. Using all the available attention matrices from all of the model’s heads, we use a linear classifier RidgeClassifier from scikit-learn^116^ (with an L2 penalty set to 1, a positivity constraint on the coefficients, and without an intercept) to classify the ground truth’s edges based on a combination of each head. The classifier converts the target values into {-1, 1} and then treats the problem as a regression task with mean squared error.

Instead of taking the classifier’s output, we directly take the average of the subset of each head associated with a non-zero coefficient in the classifier without weighting them. Thus, the classifier only serves as a means to select the heads with relevant information in predicting a ground truth of interest (see Figure 1C).

### Normalization and network interpretation

In scPRINT and scGPT, the attention matrix is normalized via the softmax function over the query (i.e. row) dimensions. This means that all row elements sum up to 1 or that the same mass flows from each network component. This rescaling is essential as it corrects that some row element scales can be much higher than others in the attention matrix. Similarly, in regularized models like GENIE3, only a small set of genes are connected for each gene in the matrix, meaning all genes have directed edges toward a small subset of genes. Thus, our interpretation is that the row elements are the targets in our network, each connected to a small subset of genes. The column elements are thus the regulators and can regulate many / most genes in the network.

For biological ground truths like MCalla et al. and gwps, which fit this assumption of highly connected regulators and sparsely regulated targets, we directly compare them to the inferred network. Table S12 and S13 show that this performs better than taking the opposite view by transposing the inferred networks.

This assumption is challenged for Omnipath, which has most of its elements connected to a sparse set of other elements (see Supp Figure S3). Due to the sparsity of connections for regulators (i.e. sources) in the ground truth network and the large number of regulators (8000+), the methods are challenged and perform much better when taking the transpose of their network and matching the regulators to the sources and sources to regulators.

### BenGRN and gene network metrics

We use the packages benGRN and GRnnData released with this manuscript to work With Gene networks and perform our benchmarks.

Our three main metrics are EPR, AUPRC, and enrichment. They all take advantage of the fact that the predictions are generated as scores over edges between nodes:

- Expected Precision Recall (EPR) is computed as the odds ratio 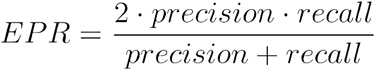 at the cutoff of the scores given by the *K* top predicted elements, where *K* is the number of positive elements in the ground truth.
- Area Under the Precision-Recall Curve (AUPRC) is the area (computed with the composite trapezoidal rule) under the curve defined by the precision (*PR = TP / (TP + FP*)) and recall (*RE = TP / (TP + FN*)) where *TP* is the number of true positives, FP is the number of false positives, and *FN* is the number of false negatives. This curve is obtained through a range of cutoffs going from 0 predicted positives to all predicted positives. Here, we compute a version of the AUPRC where the floor of the area is not given by the Precision=0 line but by the line of the prevalence of the positive class. Moreover, we do not interpolate the curve between the last recall value and the perfect recall: 1. We do this to properly compare AUPRC values across benchmarks and models. Random precision values are given in the supplementary data.
Enrichment is computed using the prerank methodology^49^, where, given an ordered set of genes, is computed by:
- 1.Summing all scores of edges of the matrix row-wise. (Target - Hub) Or
- 2.Summing all scores of edges of the matrix column-wise. (Regulators - Hub) Or
- 3.Computing the eigenvector centrality of nodes in the graph^117^ using NetworkX’s implementation. Prerank’s background comprises all the genes in the set (centrality).

Of note, we did not design an automated method for cell type enrichment. Instead, the assessment of whether or not a network is enriched for the correct cell type is done manually, identifying cell type names in the top 10 cell types listed in the enrichment results of the network.

### Other evaluation metrics

All evaluation metrics from the section scPRINT is competitive on tasks orthogonal to GN inference of the results come from the openproblems benchmark and are standards in the field.

scIB’s batch correction score is an average of the avgBatch score and the avgBio score, which are themselves averaged over many scores. Details of each value are available in our package’s notebooks.

- scIB avgBio is a combination of label-based and label-free metrics using for example: the Adjusted Rand Index (ARI)^118^ and the Normalized Mutual Information (NMI)^116^ on clusters computed from the K-Nearest Neighbor graph. Other scores are used, some using the conservation of trajectories and of the cell cycle variance, and some on the rare cell population conservation, overlap of highly variable genes (see scIB^64^), and more.
- scIB avgBatch is a similar combination of label-based and label-free metrics using for example the average connectivity across clusters of different batches: ASW^119^, the graph integration local inverse Simpson’s Index: graph iLISI^120^, as well as the the k-nearest-neighbor Batch Effect Test (kBET)^119^ and more.

Finally, we also use two metrics in our classification task:

- <u>macro-F1</u>: also called macro-average, is the average of the F1 score across each class in a multi-class task. Where the F1 score is: 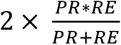.
- <u>Accuracy</u>: the accuracy is computed as 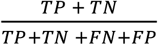

### Denoising validation test

To validate the denoising ability of scPRINT, MAGIC^60^, and KNNsmoothing2^61^, our test function, available in the scPRINT package, uses the complete set of cells in the dataset to generate the denoised expression over the 5000 most variable genes in this dataset. This is mainly done for the compute efficiency of the other models (MAGIC / KNNsmoothing2), which do not scale well with respect to the number of genes as they compute multiple PCAs over their datasets.

Before that, counts are removed from the dataset following the same procedure as done for scPRINT’s pretraining (see <u>The denoising task</u> section of the methods).

For each cell, we compare the denoised and un-denoised profiles to the “true” profile (e.g. before denoising). We compute the Spearman’s correlation over the genes initially expressed in the cell, taking the average across all cells. We do not use the unexpressed genes as we are working with a dataset with high dropout and expect that a good denoiser will set genes that are 0 in the profile with some value. We notice that this improves the score of all denoising methods and makes more sense given the data.

For the rare cell population test, we keep everything similar but compute only the Spearman correlation over a rare cell population in the dataset.

### State-of-the-art methods used in benchmarking

#### Gene network inference with ensemble of trees (GENIE3)

Developed originally for bulk transcriptional data, GENIE3 computes the regulatory network for each gene independently. It uses a random forest, a weak learner ensemble method, to predict the expression profile of each target gene from profiles of all the other genes. The weight of an interaction comes from the feature importance value of an input gene in the predictor for a target gene’s expression pattern. Aggregating these weighted interactions over all the genes yields the regulatory network. This method was the top performer in the DREAM4 in silico network challenge (multifactorial subchallenge).

GENIE3 can be seen as a generalization of correlation-based methods for infereing gene networks. Instead of looking at genes that correlate most with another gene, GENIE3 finds how to combine a set of correlated genes to get an even better “correlation”.

We also use a version of the model we call GENIE3-TF. Here, the regression is performed only using the expressed transcription factors instead of all expressed genes as input. This is the most used version of GENIE3 and is much faster.

#### Single-cell generative pretraining transformer (scGPT)

scGPT is a transformer-based model of roughly 100M parameters, pre-trained with a generative process similar to Language models. scGPT proposes to build similarity networks based on the output gene embeddings of the model but also based on its attention matrices. It computes networks as the difference between the rank-normalized version of the average attention matrix in a baseline expression profile vs a perturbed one in perturb-seq data. The attention matrix is the average of attention matrices over the heads of the last layer and over the cells given to the model.

We run scGPT following the examples given in their “Tutorial_Attention_GRN.ipynb” notebook. All runs are in our fork: “https://github.com/jkobject/scGPT“ in the “mytests/” folder. Similarly, we take the mean over cells and over the heads of the last layer. We compute softmax similarly to the attention computation but without applying the rescaling factor 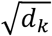. We finally drop the first element corresponding to the cell embedding token.

### Ground truth preparation

#### MCalla et al

For the MCalla et al. dataset, we downloaded the data from the supplementary datasets of their paper https://www.biorxiv.org/content/10.1101/2021.06.01.446671v2.supplementary-material. After undoing the logp1 transform, we re-generate the true count expression matrix from the normalized one by dividing the expression of each cell by the smallest value in its expression profile. This fully recovered the true counts, all values being integers. For the additional human dataset we used, we downloaded it from the gene expression atlas database https://www.ebi.ac.uk/gxa/sc/experiments/E-GEOD-36552/downloads.

We used the intersection (gold standard) ground truth dataset for both human and mouse. Converting this list of source to target genes into a directed binary network.

#### Omnipath

We generate the Omnipath network using all the interactions from the Omnipath Python package, excluding small molecules, lncRNAs, and any element without a unique HGNC symbol. We then transform it into a directed binary network of source to target.

#### Gene networks from genome-wide perturb-seq

We created a gene network from the genome-wide perturb-seq dataset using the supplementary matrix containing the results of differential expression in the dataset. This matrix represents the multiple hypothesis testing corrected p-values of a differential expression test of cells with KO of gene A compared to the baseline cell expression. This is available for all 8000+ expressed genes in the K562 cell line. We used a cutoff of 0.05 on these values to define the directed binary connection between genes.

This effectively gives a gene x gene-directed binary graph that tells if a statistically significant connection exists from the source *gene*_*A*_ to the target *gene*_*B*_ according to genome-wide perturb-seq.

For all ground truths, download, preprocessing, and extraction of the network and expression data are available in the BenGRN package.

### Details on the Benign Prostatic Hyperplasia analysis

We download our dataset from cellxgene under the reference: 574e9f9e-f8b4-41ef-bf19-89a9964fd9c7.

We preprocess the dataset using scDataloader’s preprocessing function. We generate embedding and classification using 3000 expressed genes in each cell. Similarly to pretraining, we take 3000 randomly expressed genes; if less than 3000 are expressed, we complete with randomly selected unexpressed genes. We display embeddings generated using the cell type classifier logits (see section <u>The classification task</u> in methods)

We use the Scanpy toolkit^121^ to generate our Umap plots directly from the embeddings, as well as our differential expression results and our clusters. We define the clusters using the Louvain algorithm with 10 k-nearest-neighbors and a resolution of 1. We perform denoising on 5000 genes per cell selected similarly to the embedding and classification part. We use the 4000 most variable genes in each cell type to generate our gene networks in the BPH and normal fibroblasts.

On the gene networks, we perform gene set enrichment with the Enrichr method^122^. For community detection, we use Louvain algorithm with parameter 1.5. We perform analysis only on the communities with between 200 and 20 genes. (4 and 5 in the BPH-associated fibroblasts, 3 and 4 in the normal fibroblasts)

All analysis and results are available in the *cancer_usecase_1* and *cancer_usecase_2* notebooks.

### Negative Binomial to Poisson relationship

As explained in <u>The denoising task</u> and <u>Expression decoder</u> section of the methods, in our model, we have used the ZINB as our loss, an extension of the NB distribution to zero-inflated data. Moreover, we have also used the zero-inflated Poisson - like mechanism to downsample the cell expression profiles. These are consistent because we can view the Poisson distribution as a NB without overdispersion. The relationship between *NB* and *Poisson* is given by making the dispersion term go to 0 and the inverse dispersion term *θ* → *inf*. Doing so, the term 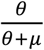 approaches 1. Thus, the PMF simplifies to:

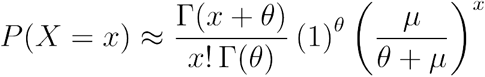

For large *θ*, we use Stirling’s approximation^123^ of the Gamma function 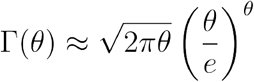 we get:

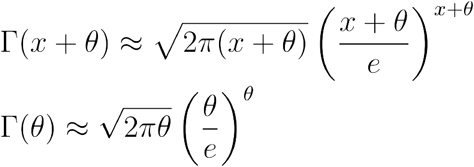

Simplifying the ratio of the Gamma functions:

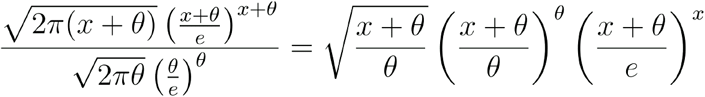

For large 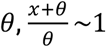, so:

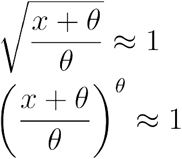

Thus, the expression simplifies to:

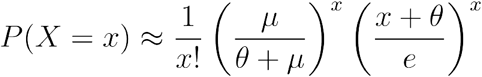

Finally, 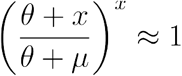 for large *θ*, so:

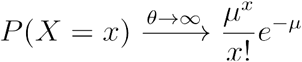

This is the PMF of the Poisson distribution with mean *μ*.

## Data availability

- model weights on: https://huggingface.co/jkobject
- pre-training logs on: https://wandb.ai/ml4ig/scprint_scale/reports/scPRINT-trainings--Vmlldzo4ODIxMjgx?accessToken=80metwx7b08hhourotpskdyaxiflq700xzmzymr6scvkp69agybt79l341tv68hp
- CellxGene datasets: https://cellxgene.cziscience.com/
- All of the other datasets used in this work can be downloaded via the helper scripts on the scPRINT, BenGRN, GRnnData and scDataLoader packages.

## Code availability

- scPRINT and notebooks to reproduce the results: https://github.com/cantinilab/scPRINT
- GrnnData package: https://github.com/cantinilab/GRnnData
- BenGRN package: https://github.com/jkobject/benGRN
- scDataLoader package: https://github.com/jkobject/scDataLoader
- scGPT and notebooks to reproduce the results: https://github.com/jkobject/scGPT/tree/main/mytests

## Acknowledgment

The project leading to this manuscript has received funding from the Inception program (Investissement d’Avenir grant ANR-16-CONV-0005) and the European Union (ERC StG, MULTIview-CELL, 101115618). We acknowledge the help of the HPC Core Facility of the Institut Pasteur and Déborah Philipps for the administrative support.

The work of G. Peyré was supported by the French government under management of Agence Nationale de la Recherche as part of the ‘Investissements d’avenir’ program, reference ANR19- P3IA-0001 (PRAIRIE 3IA Institute).

