## Supplementary information for "scPRINT: pre-training on 50 million cells allows robust gene network predictions"

### Supplementary Information: What can 50 million cells tell us about gene networks?

#### Supplementary tables

##### Table S1: List of novelties in scPRINT and comparison to scGPT and scFoundation

| features | scPRINT | scGPT | scFoundation |
|---|---|---|---|
| **classification pretraining** | v | x | x |
| **hierarchical classification** | v | x | x |
| **denoising pretraining** | v | x | v |
| **masking pretraining** | v (unused) | v | v |
| **MVC pretraining** | v (unused) | v | x |
| **bottleneck pretraining** | v | x | x |
| **large cell count GN inference** | v | x | x |
| **zero-shot classification** | v | x | x |
| **zero-shot batch correction** | v | x | x |
| **zero-shot denoising** | v | x | v |
| **genome-wide GN inference** | v | x | x |
| **large input context** | v | x | v |
| **raw count encoding** | v | x | v |
| **very large model available** | v | x | x |
| **pretraining strategy and dataset** | v | x | x |
| **low GPU/hours implementation** | v | x | x |
| **weighted random sampling pretraining** | v | x | x |
| **protein encoding** | v | x | x |
| **cross-species abilities** | v | x | x |
| **gene location encoding** | v | x | x |

| | | | |
|---|---|---|---|
| **genome-wide input context** | x | x | v |
| **xtrimogene architecture** | x | x | v |
| **Publicly available train / validate / test strategies** | v | x | x |
| **flashattention2** | v | x | x |

Comparison of the features and novelties from scPRINT compared to 2 similar published state-of-the-art methods: scGPT and scFoundation.

### Table S2: model comparison

| model name | model size | training time (hours) | training hardware | num cells | num leaf cell type | dimension (d) | layers | heads | token input size | num species | training | attention |
|---|---|---|---|---|---|---|---|---|---|---|---|---|
| **scPRINT-small** | 7M | 24 | 1xA100 | 41M (91M before QC) | 540 | 128 | 4 | 2 | 2,200 | 2 | denoising (60%) + classification + bottleneck | flashattention2 |
| **Geneformer** | ? (~20M) | 72 | 12xV100 | 30M | ? | 256 | 6 | 4 | 2,048 | 1 | masked (15%) | normal |
| **scPRINT-medium** | 20M | 72 | 1xA100 | 41M (91M before QC) | 540 | 256 | 8 | 4 | 2,200 | 2 | denoising (60%) + classification + bottleneck | flashattention2 |
| **scGPT** | 100M | ? | ? | 33M | ? | 512 | 12 | 8 | 1,200 | 1 | masked (15%) | flashattention1 |
| **scFoundation** | 100M | ? | ? | 50M | ? | 768 | 12+12 | 12+8 | 20,000 | 1 | masked (30%) + denoising | xtrimogene |
| **scPRINT** | 90M | 96 | 4xA100 | 41M (91M before QC) | 540 | 512 | 16 | 8 | 2,200 | 2 | denoising (60%) + classification + bottleneck | flashattention2 |
| **GPT2-small** | 117M | ? | ? | 300B tokens (~150M cells) | x | 768 | 12 | 12 | 1200 | x | masked (15%) | normal |
| **UCE** | 650M | 960 | 24xA100 | 36M | (~1000?) likely <500 | 1280 | 33 | 20 | 1024 | 5 | masked (20%) | normal |
| **cellFM** | 700M | ? | 32xAsce | 100M | ? | 1536 | 40 | 48 | 4096 | 1 | masked (20%) | normal + |

| | | | | | | | | | | | | |
|---|---|---|---|---|---|---|---|---|---|---|---|---|
| | | | nd910 NPUs | | | | | | | | | LORA |
| **scPRINT-vlarge** | 700M | 168 | 24xA100 | 41M (91M before QC) | 540 | 1280 | 20 | 10 | 2,200 | 2 | denoising (60%) + classification + bottleneck | flashattention2 |

Table comparing different model sizes and architectures. Comparing scPRINT to other state-of-the-art methods, as well as GPT2-small and GPT3-large models

#### Table S3: Ablation study and impact on performance across tasks

| **id** | **description** | **denoise/reco2full_vs_noisy2full** | **emb_lung/ct_class** | **emb_lung/scib** | **emb_panc/ct_class** | **emb_panc/scib** | **reconstruction loss** | **classification accuracy** | **denoising loss** | **epoch** |
|---|---|---|---|---|---|---|---|---|---|---|
| **or46096v** | small | 0.34 | 0.31 | 0.47 | 0.11 | 0.41 | 1.31 | 0.4 | 1.16 | 24 |
| **ghqf2hym** | medium | 0.12 | 0.58 | 0.55 | 0.52 | 0.51 | 1.25 | 0.33 | 1.125 | 27 |
| **7asy8qpn** | large | 0.18 | 0.69 | 0.56 | 0.52 | 0.50 | 1.23 | 0.76 | 1.109 | 21 |
| **24chcp2e** | medium-nofreeze | 0.15 | 0.45 | 0.54 | 0.52 | 0.53 | 1.25 | 0.33 | 1.115 | 23 |
| **6o76ew23** | medium-2-heads | 0.10 | 0.49 | 0.55 | 0.40 | 0.53 | 1.25 | 0.33 | 1.124 | 26 |
| **lsr3pvnf** | medium-MSE | 0.21 | 0.61 | 0.56 | 0.51 | 0.49 | 1.26 | 0.33 | 6.3 (diff) | 29 |
| **muwj73gx** | medium-MVC | 0.21 | 0.51 | 0.55 | 0.40 | 0.47 | 1.29 | 0.3 | 1.132 | 37 |
| **n8jypo8z** | medium-noPE | 0.09 | 0.71 | 0.56 | 0.35 | 0.46 | 1.27 | 0.33 | 1.31 | 23 |
| **q0fzpj5g** | medium-no-random-weighted | 0.17 | 0.51 | 0.53 | 0.19 | 0.48 | 1.26 | 0.26 | 1.118 | 27 |
| **f5e4qfkr** | medium-MLM | 0.04 | 0.53 | 0.54 | 0.39 | 0.46 | 1.26 | 0.35 | 0.999 | 23 |

The table shows the results of the ablation study on denoising, embedding with batch correction, and cell-type classification tasks. Results are displayed for the medium-size scPRINT model. Top to bottom: 1. Regular model, 2. A model trained without freezing gene embedding during pre-training, 3. a model trained with only two heads per layer instead of 4, a model with Mean Squared Error instead of the ZINB loss, 4. a version trained with scGPT's MVC methodology for the creation of the cell embedding, 5. Model trained without positional encoding for the gene's location, 6. A model trained without weighted random sampling, 7. A model trained with masked language modeling instead of denoising.

### Table S5: overlap of different GN ground truths

| comparison | precision | recall | random precision |
|---|---|---|---|
| MCalla et al. vs Omnipath | 0.0520 | 0.0074 | 0.00154 |
| MCalla et al. - T vs Omnipath | 0.0155 | 0.0022 | 0.00154 |
| gwps vs Omnipath | 0.0015 | 0.0219 | 0.00129 |
| gwps -T vs Omnipath | 0.0030 | 0.0426 | 0.00129 |

Comparison of the overlap, expressed as precision and recall, of the three different ground truth networks used: MCalla, Omnipath, and gwps.

### Table S6: Omnipath benchmark results on the genome-wide perturb-seq dataset

| tool | EPR | AUPRC | TF target enr. | TF_enr | TF_only | ct_pred | RAND precision |
|---|---|---|---|---|---|---|---|
| **genie3** | 4.68 | 0.00188 | 17.9 | TRUE | FALSE | FALSE | 0.00163 |
| **scGPT** | 0.99 | 0.00208 | 14.0 | TRUE | FALSE | FALSE | 0.00163 |
| **scPRINT** | 2.81 | 0.00170 | 8.6 | TRUE | FALSE | FALSE | 0.00161 |
| **scPRINT-omni** | 4.70 | 0.00189 | 3.4 | TRUE | FALSE | FALSE | 0.00161 |
| **scPRINT-self** | 1.61 | 0.00190 | 5.0 | TRUE | FALSE | FALSE | 0.00161 |

Omnipath network overlap (EPR, AUPRC), as well as transcription factor enrichment, TF target enrichment, and cell type marker enrichment for gene networks generated by the different tools on the genome-wide perturb seq K562 cells at steady state (no perturbations)

### Table S7: Omnipath benchmark results on the MCalla et al. datasets

| tool | dataset | EPR | AUPRC | TF target enr. | TF enr. | cell type enr. |
|---|---|---|---|---|---|---|
| **genie3** | **Han et al.** | 1.51 | 0.00016 | 11.3 | FALSE | TRUE |
| **genie3** | **Yan et al.** | 1.74 | 0.00020 | 0.0 | FALSE | TRUE |
| **scGPT** | **Han et al.** | 0.89 | 0.00016 | 17.0 | TRUE | FALSE |
| **scGPT** | **Yan et al.** | 0.16 | 0.00007 | 20.0 | FALSE | FALSE |
| **scPRINT** | **Han et al.** | 2.03 | 0.00019 | 23.6 | TRUE | FALSE |
| **scPRINT** | **Yan et al.** | 1.76 | 0.00026 | 31.1 | FALSE | TRUE |
| **scPRINT-omni** | **Han et al.** | 5.12 | 0.00004 | 3.6 | TRUE | FALSE |

| | | | | | | |
|---|---|---|---|---|---|---|
| **scPRINT-omni** | **Yan et al.** | 3.35 | 0.00019 | 13.3 | FALSE | TRUE |
| **scPRINT-self** | **Han et al.** | 0.94 | 0.00030 | 30.9 | TRUE | TRUE |
| **scPRINT-self** | **Yan et al.** | 0.57 | -0.00004 | 6.7 | TRUE | TRUE |

Omnipath network overlap (EPR, AUPRC), as well as transcription factor enrichment, TF target enrichment and cell type marker enrichment for gene networks generated by the different tools on the 2 human embryonic stem cell datasets used in scPRINT outperforms GENIE3 and scGPT on cell type-specific ground truths.

### Table S8: Denoising results per datasets

| tools | denoising (+%) correlation. colon | denoising (+%) correlation. retina | denoising (+%) correlation. ciliary body | denoising (+%) correlation (low cell count: 30). colon | denoising (+%) correlation (low cell count: 30). retina | denoising (+%) correlation (low cell count: 30). ciliary body | average denoising (+%) correlation | average denoising (+%) correlation (rare cell type) |
|---|---|---|---|---|---|---|---|---|
| **random scPRINT model** | -16.0 | X | X | -16.0 | X | X | -16.0 | -16.0 |
| **scPRINT-large** | 19.1 | 33.9 | 17.1 | 22.5 | 26.6 | 16.6 | 23.4 | 21.9 |
| **KNNsmoothing2** | 21.0 | 34.9 | 21.6 | 17.0 | 32.0 | 13.4 | 25.8 | 20.8 |
| **magic** | 29.3 | 34.6 | 22.7 | 16.8 | 24.4 | 4.6 | 28.9 | 15.3 |
| **magic (low cell dataset)** | X | X | X | 11.3 | 14.0 | 13.0 | X | 12.8 |

This table shows the detail of the denoising results for each of the three datasets for scPRINT-large, KNNsmoothing2, MAGIC, and MAGIC run on only the small cell type cluster. “Random scPRINT model” is the performance of an untrained scPRINT model.

### Table S9: highlighted b-cell cluster genes in the BPH study

| gene | link | in cancer | in b cell | analysis |
|---|---|---|---|---|
| **MBNL2** | https://www.nature.com/articles/s41467-023-44126-w | prostate cancer | high expr in immune tissues | BPH B-cell to normal B-cell diff. expr. |
| **MAGOH** | https://www.ncbi.nlm.nih.gov/pmc/articles/PMC9738831/ | cancer | high expr in immune tissues | |
| **RANBP2** | https://www.nature.com/articles/leu2012286. https://www.genecards.org/cgi-bin/card | B-cell lymphoma | b cell validated | |

| | | | | |
|---|---|---|---|---|
| | disp.pl?gene=RANBP2 | | | |
| **CLIC4** | https://www.nature.com/articles/s41420-022-01003-7 | prostate cancer | high expr in immune tissues | |
| **BAG5** | https://www.ncbi.nlm.nih.gov/pmc/articles/PMC3598994/ | prostate cancer | b cell in cancer | |
| **NR4A1** | https://www.ncbi.nlm.nih.gov/pmc/articles/PMC9424640/ https://www.ncbi.nlm.nih.gov/pmc/articles/PMC8081071/ | prostate cancer | b cell validated | |
| **BAZ2A** | https://www.nature.com/articles/s41598-024-56073-7 | prostate cancer | b cell validated | |
| **ZBTB16** | https://www.pnas.org/doi/full/10.1073/pnas.0703872104 https://www.ncbi.nlm.nih.gov/pmc/articles/PMC5642638/ | tumor suppressor in prostate cancer | b cell validated | |
| **TAP1** | https://bmccancer.biomedcentral.com/articles/10.1186/s12885-023-10527-9 https://www.ncbi.nlm.nih.gov/pmc/articles/PMC5674960/ | cancer | b cell validated | |
| **TAS2R19** | https://v19.proteinatlas.org/ENSG00000212124-TAS2R19/tissue/B-cells | | b cell validated | |
| **PRDM7** | https://pubmed.ncbi.nlm.nih.gov/27129774/ | cancer | | |
| **TSEN54** | https://www.ncbi.nlm.nih.gov/pmc/articles/PMC10120902/ | cancer | b cell in cancer | |
| **EHMT2** | https://www.frontiersin.org/journals/immunology/articles/10.3389/fimmu.2017.00429/full | | b cell validated | |
| **ERICH6B** | https://platform.opentargets.org/target/ENSG00000163645/associations | cancer | | |
| **IL10RB** | https://pubmed.ncbi.nlm.nih.gov/37144812/ | cancer | b cell in cancer | BPH B-cell to normal B-cell diff. expr. post denoising |

Table of the highlighted genes in the differential expression analysis in BPH vs normal B-cells together with their annotation on their relation to cancer and to b-cells, with sources.

#### Table S10: hub and differential hub genes in the fibroblast GN of the BPH study

| TOP 15 hubs in BPH fibroblasts GN | TOP 15 hubs in normal fibroblasts GN | TOP 15 differential hubs in BPH fibroblasts vs normal | TOP 15 eigenvector_centrality differential hubs in BPH fibroblasts vs normal |
|---|---|---|---|
| HSPA1A | S100A6 | HLA-A | CD99 |
| MT2A | TGIF2-RAB5IF | MT2A | HLA-A |
| CREM | MIF | ATP6V0C | HSPA1A |
| TGIF2-RAB5IF | DNAJB9 | DEFA1 | LUM |
| HSPE1 | IGFBP7 | EIF4A1 | ATP6V0C |
| CALD1 | APOD | HSPA1A | CD99 |
| SPOCK3 | BRME1 | LUM | EIF4A1 |
| HLA-A | SPARCL1 | SPOCK3 | PAGE4 |
| SPARCL1 | TIMP1 | nan-99 | RYR2 |
| RBP1 | DCN | CD99 | SERPINF1 |
| C1S | C1S | CPE | C1R |
| BRME1-1 | MGP | THBS1 | COL6A2 |
| FABP4 | nan-270 | LGALS1 | HNRNPA0 |
| nan-99 | SLC25A6 | PYDC2 | SERPING1 |
| LUM | BLOC1S5-TXNDC5 | SERPING1 | SERPINA3 |

List of the Top-15 elements in different GN analyses. Genes in yellow in the last columns are the new ones found with eigenvector centrality compared to the 3rd columns.

#### Table S11: number of elements predicted per class

| | |
|---|---|
| ethnicity | 21 |
| sex | 2 |
| organism | 2 |

| | |
|---|---|
| cell type | 424 |
| disease | 62 |
| assay | 26 |

Number of labels predicted by the model for each class. We use hierarchical classification for cell type, disease, assay, and ethnicity.

### Table S12: full MCalla et al. results

| tool | name | EPR | AUPRC | RAND | TF_only |
|---|---|---|---|---|---|
| **genie3 (T)** | **Han et al.** | 1.14 | 0.0288 | 0.0274 | TRUE |
| **genie3 (T)** | **Han et al. (ChIP)** | 1.34 | 0.2843 | 0.2537 | TRUE |
| **genie3 (T)** | **Han et al. (KO)** | 0.91 | 0.0696 | 0.0766 | TRUE |
| **genie3 (T)** | **Han et al.** | 1.47 | 0.0955 | 0.0766 | FALSE |
| **genie3 (T)** | **Han et al. (ChIP)** | 0.96 | 0.2429 | 0.2537 | FALSE |
| **genie3 (T)** | **Han et al. (KO)** | 1.47 | 0.0955 | 0.0766 | FALSE |
| **genie3 (T)** | **Yan et al.** | 1.36 | 0.0374 | 0.0318 | TRUE |
| **genie3 (T)** | **Yan et al.** | 1.35 | 0.0366 | 0.0318 | FALSE |
| **genie3 (T)** | **Tran et al.** | 0.43 | 0.0306 | 0.0405 | TRUE |
| **genie3 (T)** | **Tran et al. (ChIP)** | 1.49 | 0.2298 | 0.1989 | TRUE |
| **genie3 (T)** | **Tran et al. (KO)** | 0.99 | 0.0828 | 0.0796 | TRUE |
| **genie3 (T)** | **Tran et al.** | 0.75 | 0.0698 | 0.0796 | FALSE |
| **genie3 (T)** | **Tran et al. (ChIP)** | 1.16 | 0.2074 | 0.1989 | FALSE |
| **genie3 (T)** | **Tran et al. (KO)** | 0.75 | 0.0698 | 0.0796 | FALSE |
| **genie3 (T)** | **Zhao et al.** | 1.14 | 0.0565 | 0.0531 | TRUE |
| **genie3 (T)** | **Zhao et al.** | 1.28 | 0.0573 | 0.0531 | FALSE |
| **scGPT (T)** | **Han et al.** | 1.22 | 0.0316 | 0.0274 | FALSE |
| **scGPT (T)** | **Han et al. (ChIP)** | 0.95 | 0.2486 | 0.2537 | FALSE |
| **scGPT (T)** | **Han et al. (KO)** | 1.49 | 0.0957 | 0.0766 | FALSE |
| **scGPT (T)** | **Yan et al.** | 1.02 | 0.0325 | 0.0322 | FALSE |
| **scPRINT-mean (T)** | **Han et al.** | 1.18 | 0.0409 | 0.0390 | FALSE |
| **scPRINT-mean (T)** | **Han et al. (ChIP)** | 1.06 | 0.2711 | 0.2642 | FALSE |

| | | | | | |
|---|---|---|---|---|---|
| **scPRINT-mean (T)** | **Han et al. (KO)** | 1.10 | 0.0863 | 0.0827 | FALSE |
| **scPRINT-mean (T)** | **Yan et al.** | 0.97 | 0.0275 | 0.0270 | FALSE |
| **scPRINT-mean (T)** | **Tran et al.** | 1.33 | 0.0442 | 0.0402 | FALSE |
| **scPRINT-mean (T)** | **Tran et al. (ChIP)** | 1.09 | 0.2128 | 0.2011 | FALSE |
| **scPRINT-mean (T)** | **Tran et al. (KO)** | 1.07 | 0.0870 | 0.0858 | FALSE |
| **scPRINT-mean (T)** | **Zhao et al.** | 1.20 | 0.0445 | 0.0392 | FALSE |
| **scPRINT-omni (T)** | **Han et al.** | 0.53 | 0.0357 | 0.0390 | FALSE |
| **scPRINT-omni (T)** | **Han et al. (ChIP)** | 0.77 | 0.2443 | 0.2642 | FALSE |
| **scPRINT-omni (T)** | **Han et al. (KO)** | 0.73 | 0.0776 | 0.0827 | FALSE |
| **scPRINT-omni (T)** | **Yan et al.** | 0.46 | 0.0260 | 0.0270 | FALSE |
| **scPRINT-omni (T)** | **Tran et al.** | 0.65 | 0.0357 | 0.0402 | FALSE |
| **scPRINT-omni (T)** | **Tran et al. (ChIP)** | 0.61 | 0.1742 | 0.2011 | FALSE |
| **scPRINT-omni (T)** | **Tran et al. (KO)** | 0.88 | 0.0825 | 0.0858 | FALSE |
| **scPRINT-omni (T)** | **Zhao et al.** | 0.47 | 0.0324 | 0.0392 | FALSE |
| **scPRINT-self (T)** | **Han et al.** | 1.29 | 0.0928 | 0.0827 | FALSE |
| **scPRINT-self (T)** | **Han et al. (ChIP)** | 0.96 | 0.2593 | 0.2642 | FALSE |
| **scPRINT-self (T)** | **Han et al. (KO)** | 1.29 | 0.0928 | 0.0827 | FALSE |
| **scPRINT-self (T)** | **Yan et al.** | 1.45 | 0.0278 | 0.0270 | FALSE |
| **scPRINT-self (T)** | **Tran et al.** | 1.33 | 0.0409 | 0.0402 | FALSE |
| **scPRINT-self (T)** | **Tran et al. (ChIP)** | 1.07 | 0.2012 | 0.2011 | FALSE |
| **scPRINT-self (T)** | **Tran et al. (KO)** | 1.09 | 0.0875 | 0.0858 | FALSE |
| **scPRINT-self (T)** | **Zhao et al.** | 1.24 | 0.0506 | 0.0467 | FALSE |

| | | | | | |
|---|---|---|---|---|---|
| **genie3** | **Han et al.** | 1.44 | 0.0292 | 0.0274 | FALSE |
| **genie3** | **Han et al. (ChIP)** | 1.21 | 0.2766 | 0.2537 | FALSE |
| **genie3** | **Han et al. (KO)** | 1.12 | 0.0775 | 0.0766 | FALSE |
| **genie3** | **Han et al.** | 1.12 | 0.0775 | 0.0766 | TRUE |
| **genie3** | **Han et al. (ChIP)** | 0.78 | 0.2621 | 0.2537 | TRUE |
| **genie3** | **Han et al. (KO)** | 1.12 | 0.0775 | 0.0766 | TRUE |
| **genie3** | **Yan et al.** | 2.17 | 0.0282 | 0.0247 | TRUE |
| **genie3** | **Yan et al.** | 1.09 | 0.0286 | 0.0247 | FALSE |
| **genie3** | **Tran et al.** | 1.22 | 0.0424 | 0.0405 | TRUE |
| **genie3** | **Tran et al. (ChIP)** | 1.48 | 0.2291 | 0.1989 | TRUE |
| **genie3** | **Tran et al. (KO)** | 0.98 | 0.0824 | 0.0796 | TRUE |
| **genie3** | **Tran et al.** | 0.97 | 0.0737 | 0.0796 | FALSE |
| **genie3** | **Tran et al. (ChIP)** | 0.95 | 0.1972 | 0.1989 | FALSE |
| **genie3** | **Tran et al. (KO)** | 0.97 | 0.0737 | 0.0796 | FALSE |
| **genie3** | **Zhao et al.** | 1.62 | 0.0577 | 0.0531 | TRUE |
| **genie3** | **Zhao et al.** | 1.06 | 0.0579 | 0.0531 | FALSE |
| **scGPT** | **Han et al.** | 0.39 | 0.0218 | 0.0274 | FALSE |
| **scGPT** | **Han et al. (ChIP)** | 1.25 | 0.2444 | 0.2537 | FALSE |
| **scGPT** | **Han et al. (KO)** | 0.20 | 0.0626 | 0.0766 | FALSE |
| **scGPT** | **Yan et al.** | 0.01 | 0.0244 | 0.0321 | FALSE |
| **scPRINT-mean** | **Han et al.** | 2.98 | 0.0695 | 0.0390 | FALSE |
| **scPRINT-mean** | **Han et al. (ChIP)** | 1.15 | 0.2946 | 0.2642 | FALSE |
| **scPRINT-mean** | **Han et al. (KO)** | 1.94 | 0.0964 | 0.0827 | FALSE |
| **scPRINT-mean** | **Yan et al.** | 0.03 | 0.0244 | 0.0270 | FALSE |
| **scPRINT-mean** | **Tran et al.** | 2.98 | 0.0507 | 0.0402 | FALSE |
| **scPRINT-mean** | **Tran et al. (ChIP)** | 1.15 | 0.2221 | 0.2011 | FALSE |
| **scPRINT-mean** | **Tran et al. (KO)** | 1.24 | 0.0912 | 0.0858 | FALSE |
| **scPRINT-mean** | **Zhao et al.** | 0.24 | 0.0429 | 0.0392 | FALSE |
| **scPRINT-omni** | **Han et al.** | 4.78 | 0.0510 | 0.0390 | FALSE |
| **scPRINT-omni** | **Han et al. (ChIP)** | 0.77 | 0.2443 | 0.2642 | FALSE |

| | | | | | |
|---|---|---|---|---|---|
| **scPRINT-omni** | **Han et al. (KO)** | 0.73 | 0.0776 | 0.0827 | FALSE |
| **scPRINT-omni** | **Yan et al.** | 0.14 | 0.0221 | 0.0270 | FALSE |
| **scPRINT-omni** | **Tran et al.** | 0.35 | 0.0351 | 0.0402 | FALSE |
| **scPRINT-omni** | **Tran et al. (ChIP)** | 0.65 | 0.1729 | 0.2011 | FALSE |
| **scPRINT-omni** | **Tran et al. (KO)** | 1.36 | 0.0921 | 0.0858 | FALSE |
| **scPRINT-omni** | **Zhao et al.** | 0.48 | 0.0347 | 0.0392 | FALSE |
| **scPRINT-self** | **Han et al.** | 3.79 | 0.1753 | 0.0827 | FALSE |
| **scPRINT-self** | **Han et al. (ChIP)** | 0.92 | 0.2601 | 0.2642 | FALSE |
| **scPRINT-self** | **Han et al. (KO)** | 3.79 | 0.1753 | 0.0827 | FALSE |
| **scPRINT-self** | **Yan et al.** | 2.58 | 0.0415 | 0.0270 | FALSE |
| **scPRINT-self** | **Tran et al.** | 7.78 | 0.1157 | 0.0402 | FALSE |
| **scPRINT-self** | **Tran et al. (ChIP)** | 1.43 | 0.2334 | 0.2011 | FALSE |
| **scPRINT-self** | **Tran et al. (KO)** | 2.08 | 0.1138 | 0.0858 | FALSE |
| **scPRINT-self** | **Zhao et al.** | 2.79 | 0.0774 | 0.0392 | FALSE |

Table showing the results of scPRINT-self, scPRINT-omni, scPRINT-mean, GENIE3, and scGPT on the MCalla et al. benchmark, also given in Figure 3B. We add the results when transposing the gene networks inferred by each model (-T)

### Table S13: full gwps results

| **tools** | **EPR** | **AUPRC** | **RAND** | **TF_only** | **AUPRC (before correction** |
|---|---|---|---|---|---|
| **genie3 - T** | 3.51 | 0.0148 | 0.0207 | FALSE | 0.0355 |
| **genie3 - TF - T** | 1.08 | 0.0001 | 0.0207 | TRUE | 0.0208 |
| **scGPT - T** | 0.43 | -0.0028 | 0.0200 | FALSE | 0.0173 |
| **scPRINT - T** | 1.10 | 0.0029 | 0.0200 | FALSE | 0.0229 |
| **scPRINT-omni - T** | 0.74 | 0.0007 | 0.0200 | FALSE | 0.0207 |
| **scPRINT-self - T** | 4.83 | 0.0230 | 0.0200 | FALSE | 0.0430 |
| **genie3** | 3.54 | 0.0149 | 0.0207 | FALSE | 0.0357 |
| **genie3 - TF** | 1.07 | 0.0001 | 0.0207 | TRUE | 0.0208 |
| **scGPT** | 2.03 | 0.0064 | 0.0200 | FALSE | 0.0264 |
| **scPRINT** | 2.71 | 0.0106 | 0.0200 | FALSE | 0.0306 |

| | | | | | |
|---|---|---|---|---|---|
| **scPRINT-omni** | 1.21 | 0.0009 | 0.0200 | FALSE | 0.0209 |
| **scPRINT-self** | 3.77 | 0.0169 | 0.0200 | FALSE | 0.0369 |

Table showing the results of scPRINT-self, scPRINT-omni, scPRINT-mean, GENIE3, and scGPT on the gwps benchmark, also given in Figure 3C. We add the results when transposing the gene networks inferred by each model (-T)

### Supplementary figures

#### FIG S1: visualization of human gene embedding from ESM2

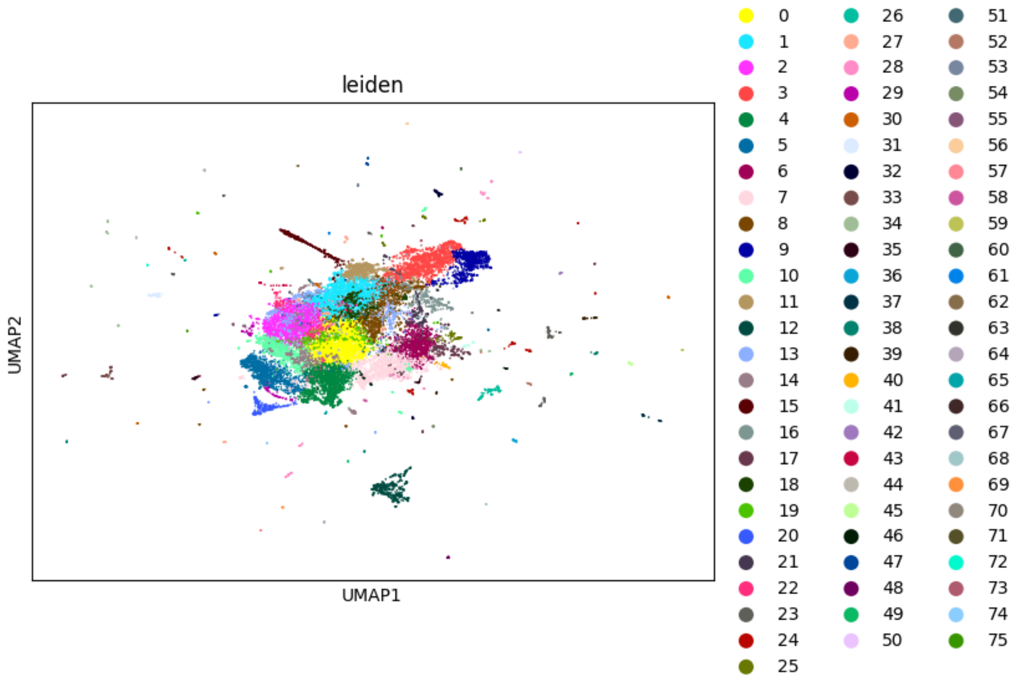

Umap of the ESM2 protein embeddings for the most common protein of all protein coding genes in Ensembl. The PCA variance ratio is 0.856 for the top 50 principal components. We color it using the louvain clustering of the embedding.

### FIG S2: Gene network inference comparison with Omnipath per datasets

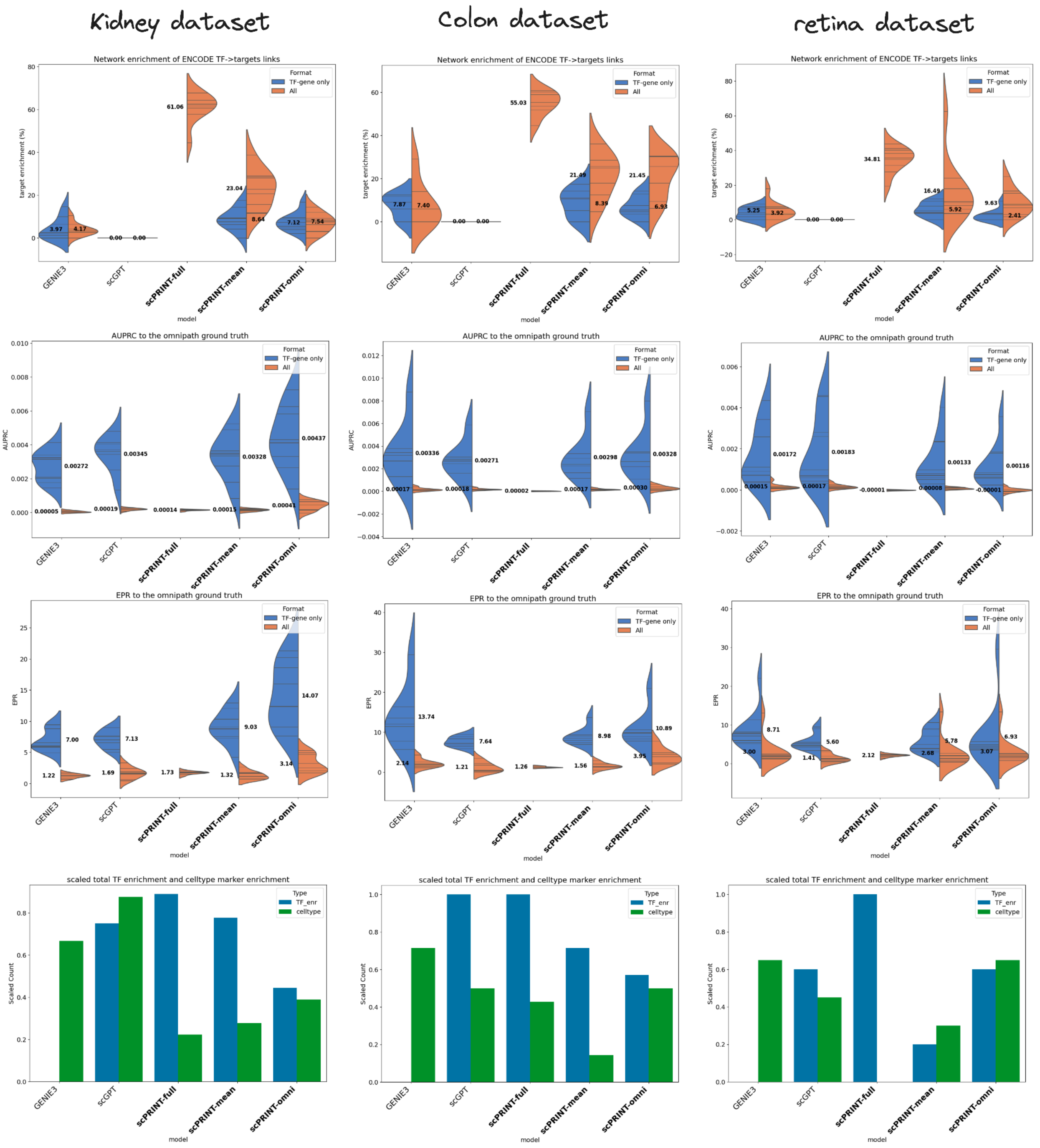

The same plots as in Figure 2B, C, and D showing the Omnipath and enrichment results per dataset for each of the 3 datasets used.

### FIG S3: Distribution of connection amongst the three ground truths

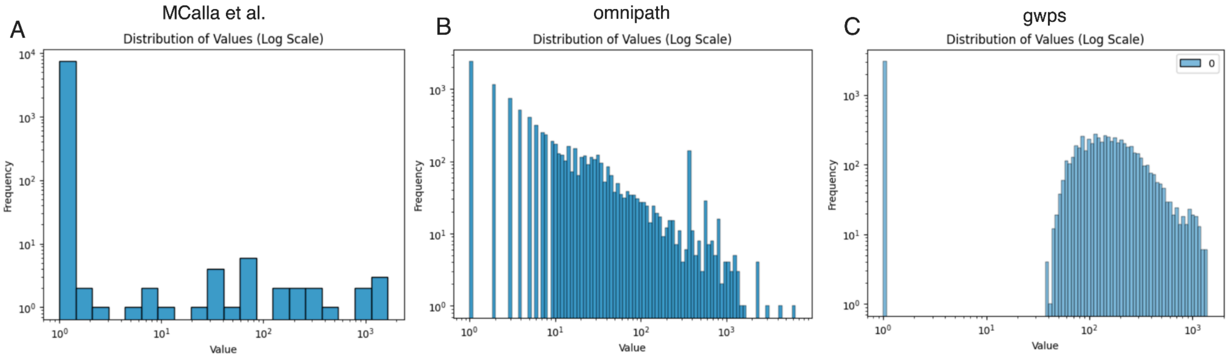

(a) Barplot of the distribution of the number of connections per edge in the MCalla human ground truth network. Most connections are 0, and there is a roughly uniform distribution of connections otherwise. This means most connections belong to the half a dozen most connected edges. (b) Barplot of the distribution of the number of connections per edge in the Omnipath ground truth network. We can see an almost linear relationship on the log-log scale, suggesting a power law distribution. (c)  Barplot of the distribution of the number of connections per edge in the genome-wide perturb-seq ground truth network. We can see a very different distribution where only a few genes have little differentially expressed genes post-knock-out, and this trend increases until reaching around 200 connections. Then, it diminishes in what might be a power law. However, some of it is likely caused by the differential expression method and noise in the scRNAseq methodology.

### FIG S4: Full denoising results

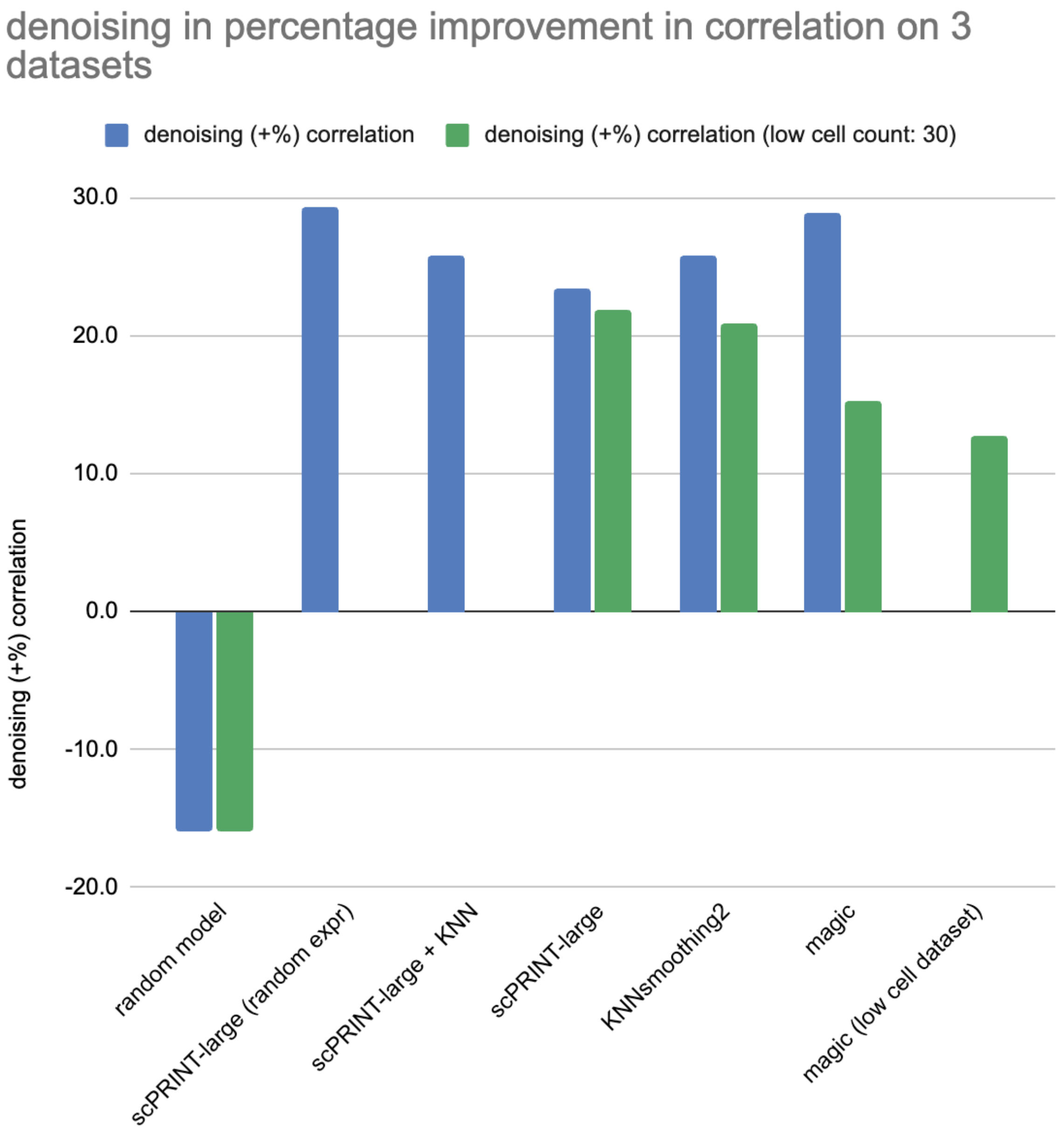

Denoising scores, similar to Figure 4A, but over more tools. “Random model” means a scPRINT model without pre-training. “Random expr” means that scPRINT was using a set of 3000 genes in a similar way as done in pre-training: Taking random expressed genes completed with random unexpressed genes if less than 3000 genes are expressed in the cell. “low cell dataset” means that MAGIC was only using the rare cell population for the dataset as presented in section Denoising validation test of the methods.

### FIG S5: Full cell type classification metrics

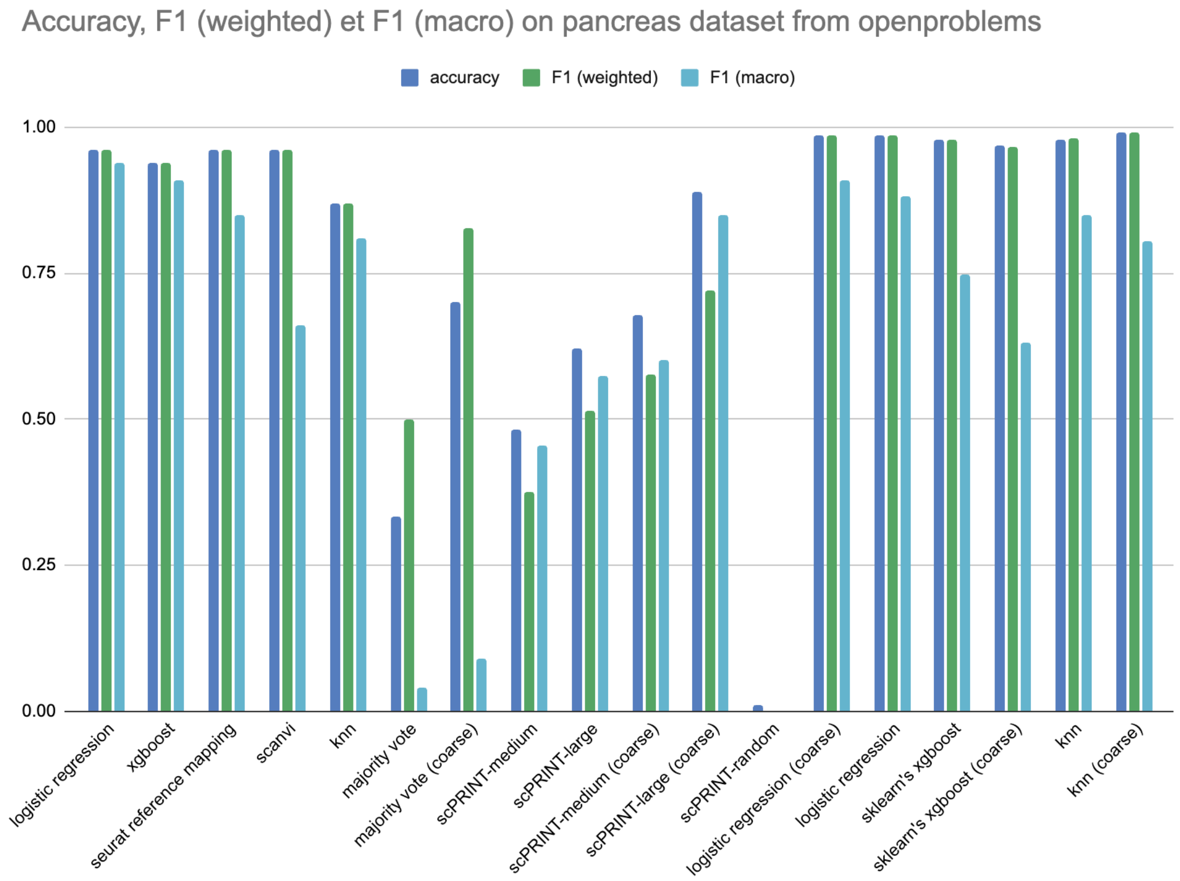

Cell type classification scores over the kidney test dataset of openproblems. Same as Figure 4B but over more tools. Coarse represents the coarse labels defined in the section [scPRINT is competitive on tasks orthogonal to GN inference] of the results.

### FIG S6: Full scIB batch correction scores

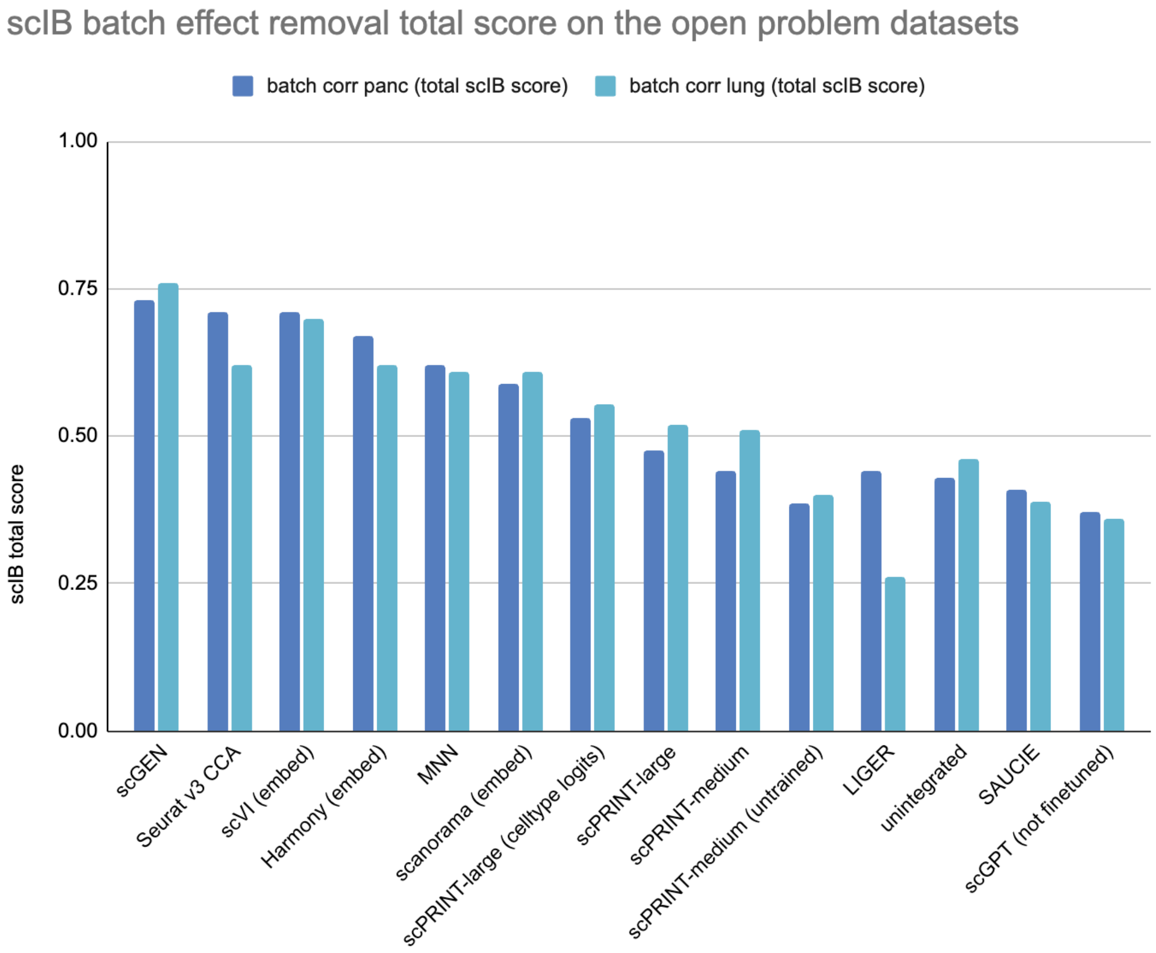

scIB benchmarking scores, averaged for the kidney and lung openproblems test datasets. Same as Figure 4C but over more tools. Cell type logits mean that the logits of the cell type classifier have been used as cell embeddings instead of the cell type embedding itself.

#### FIG S7: Full avgBio scores

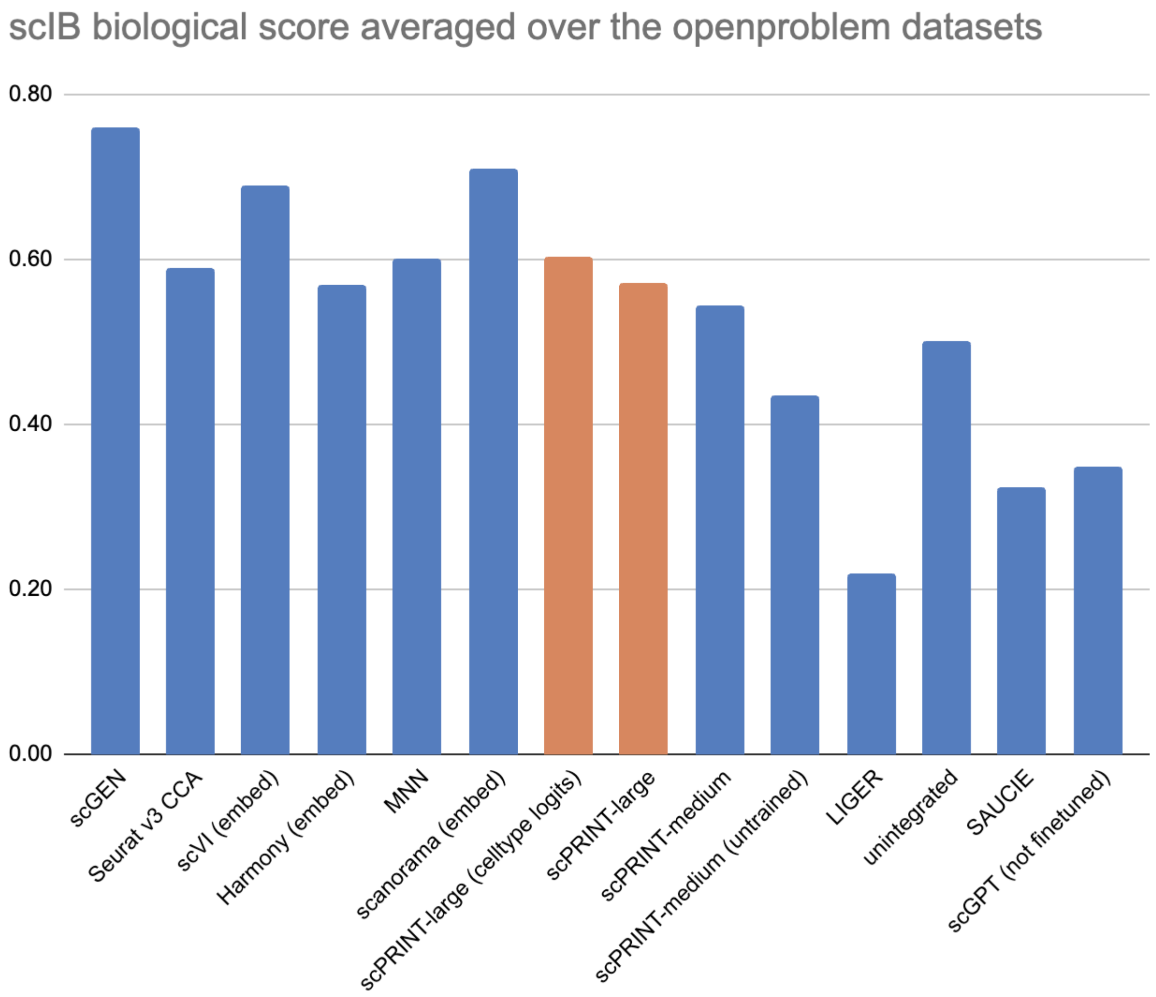

The average Biological score of the scIB benchmark averaged over the kidney and lung openproblems test datasets. Same as Figure 4D but over more tools. Cell type logits mean that the logits of the cell type classifier have been used as cell embeddings instead of the cell type embedding itself.

### FIG S8: In-depth view of the BPH dataset and its scPRINT-predicted annotations

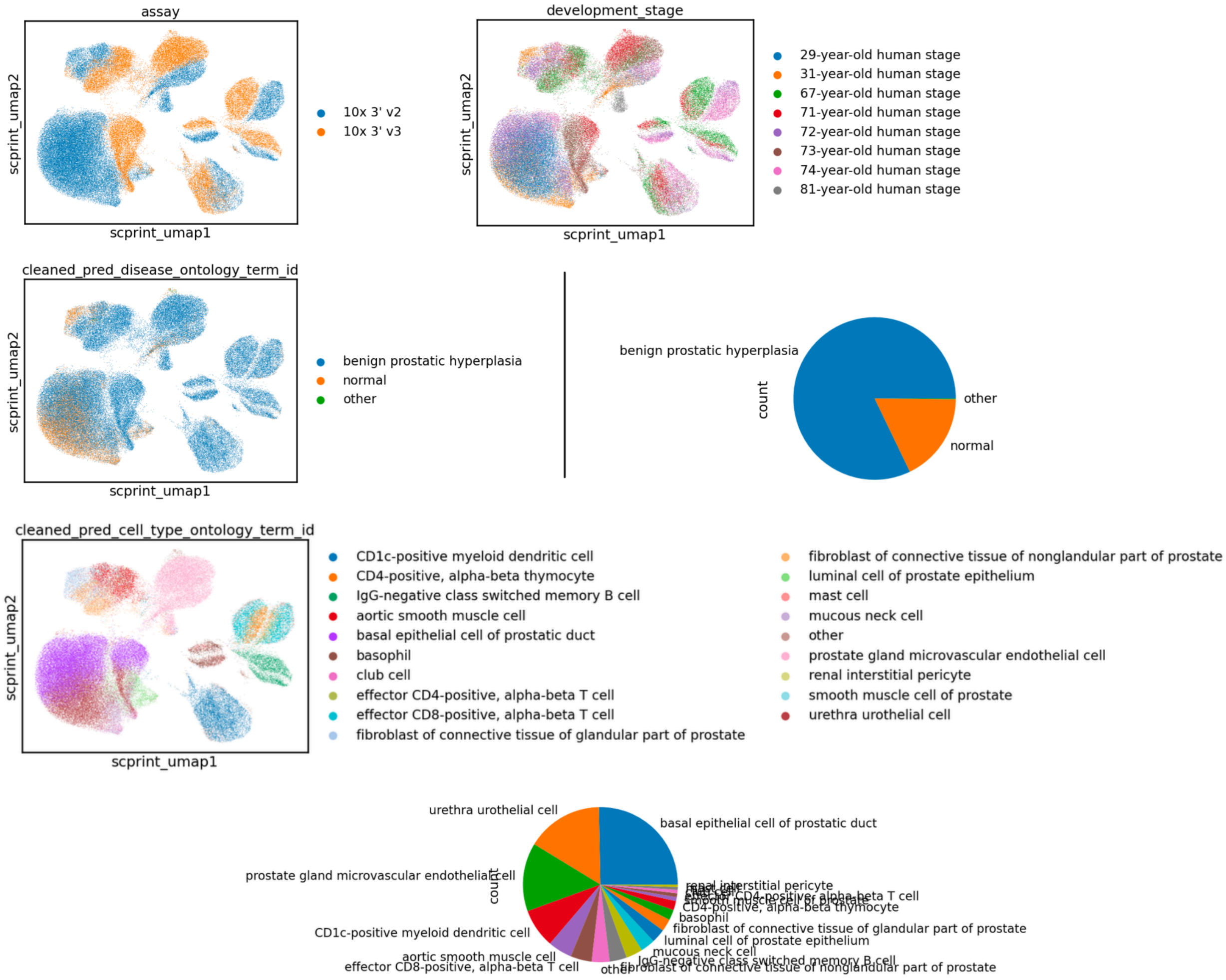

Detailed view of the assay, development stage, scPRINT-predicted diseases, and scPRINT-predicted cell types. Predicted diseases and cell types have been “cleaned” following the strategy presented in Figure 5. We also add pie charts of the relative abundance of each predicted label.

**FIG S9: differential expression analysis of the B-cell cluster vs the rest of the cells in the BPH dataset**

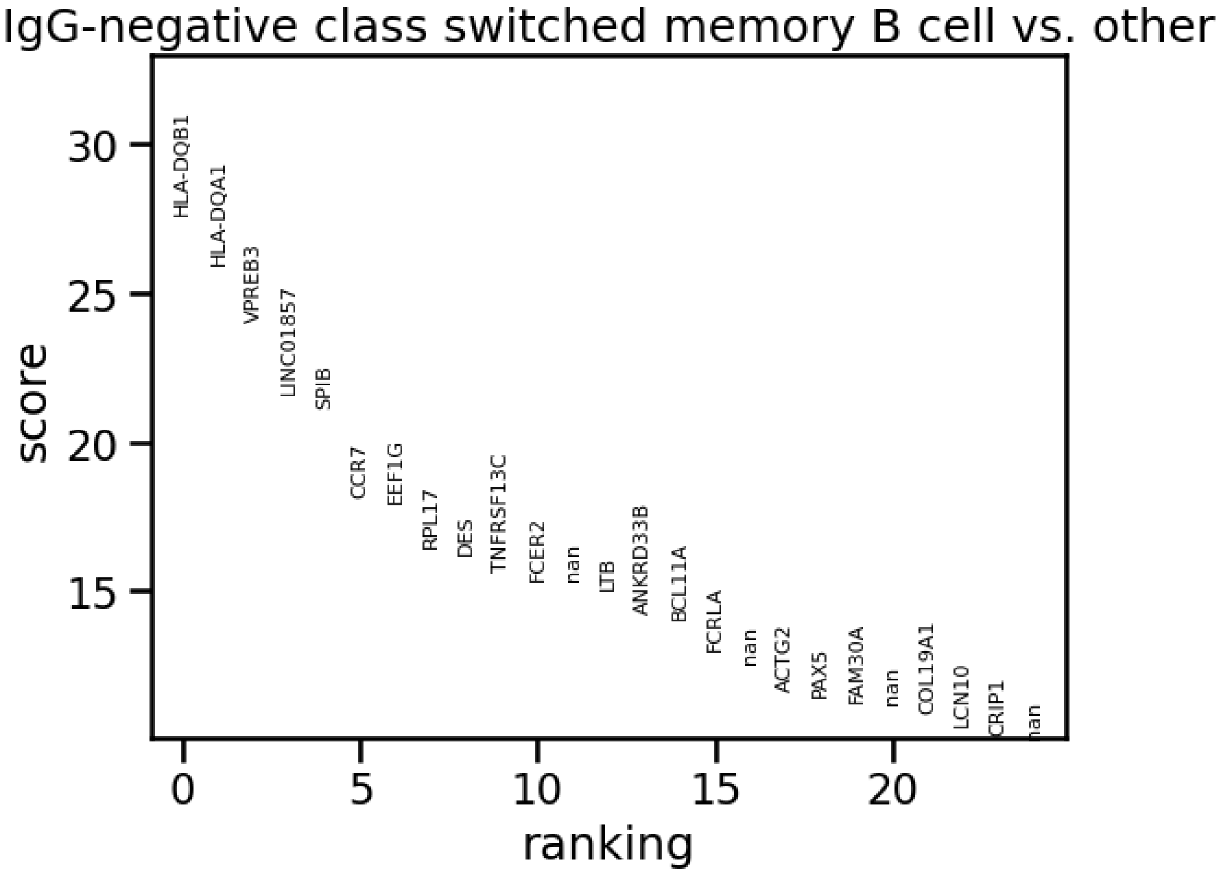

Top genes of the differential expression analysis of the scPRINT inferred B-cell cluster in vs the rest of the cells in the BPH dataset.

### FIG S10: gene enrichment comparison in the PAGE4 GN

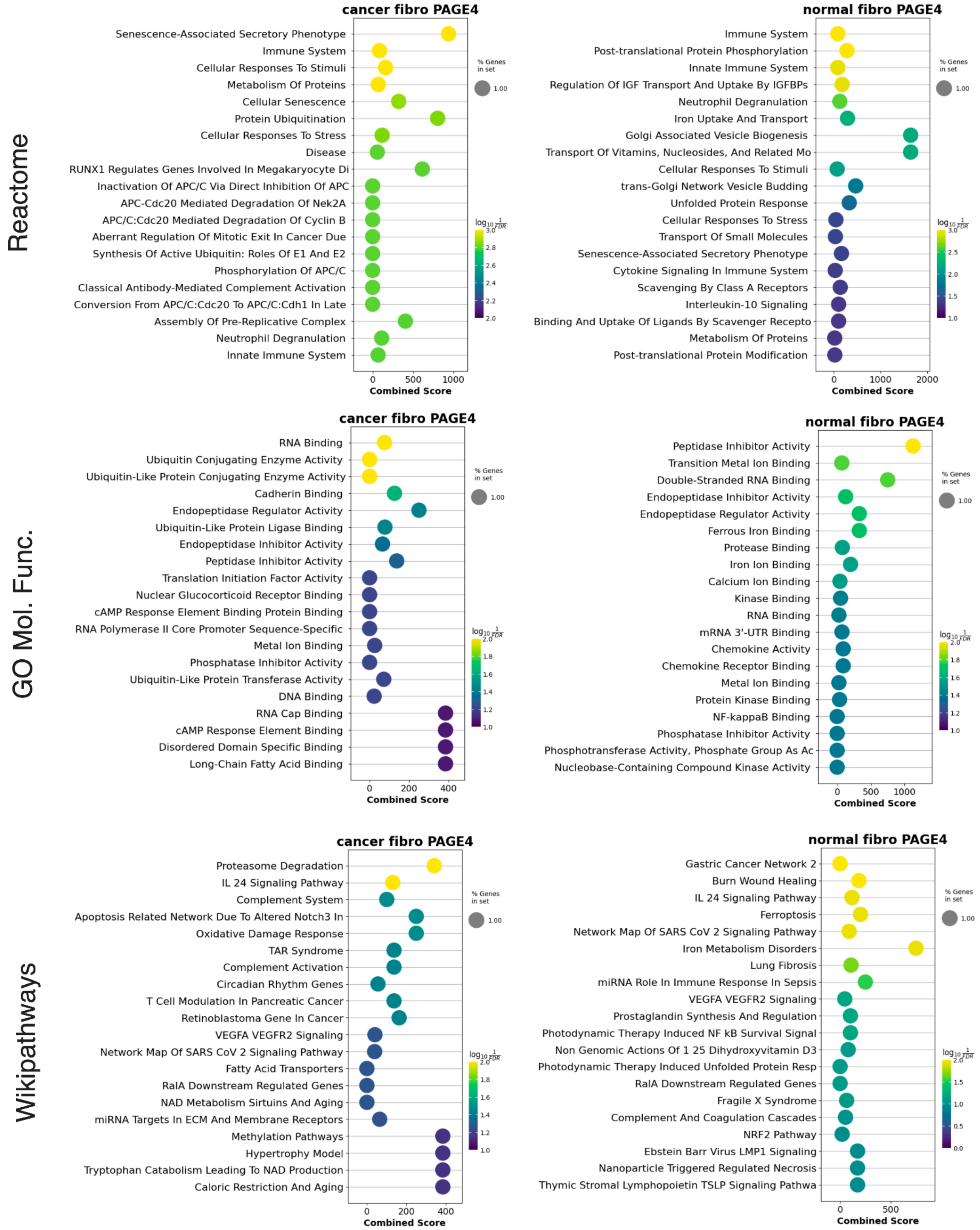

Comparison of the top 20 most enriched terms in Wikipathways, GO molecular function, and Reactome for the 40 most connected genes to PAGE4 in both BPH-associated and normal fibroblast GNs inferred by scPRINT

### FIG S11: Gene Network enrichment comparison between the BPH and normal fibroblast on their Louvain communities

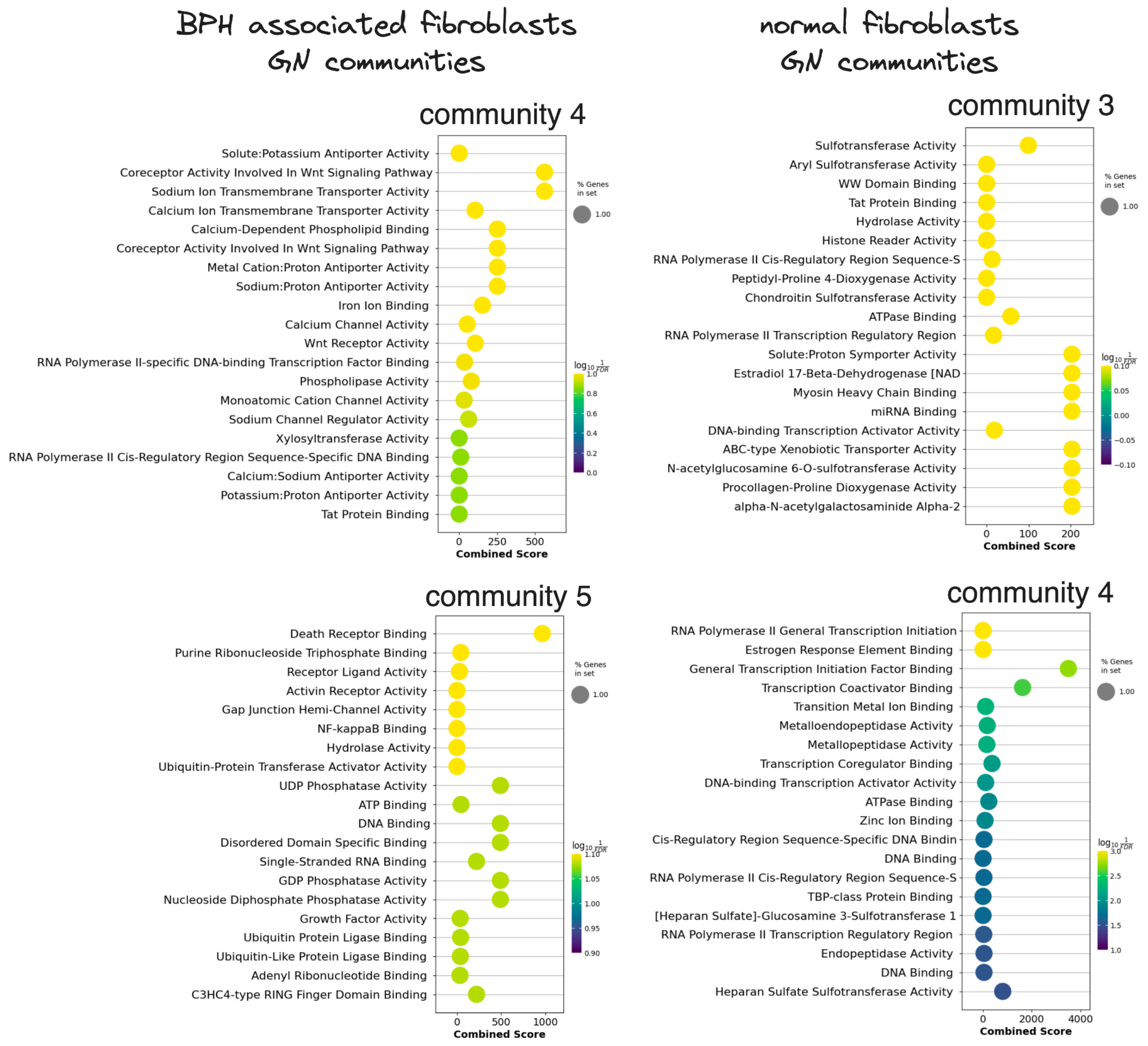

Dotplot of the top 20 GO Molecular function gene sets enriched in the Louvain communities of the BPH and normal fibroblast's Gene Networks.

#### FIG S12: Graphical Model

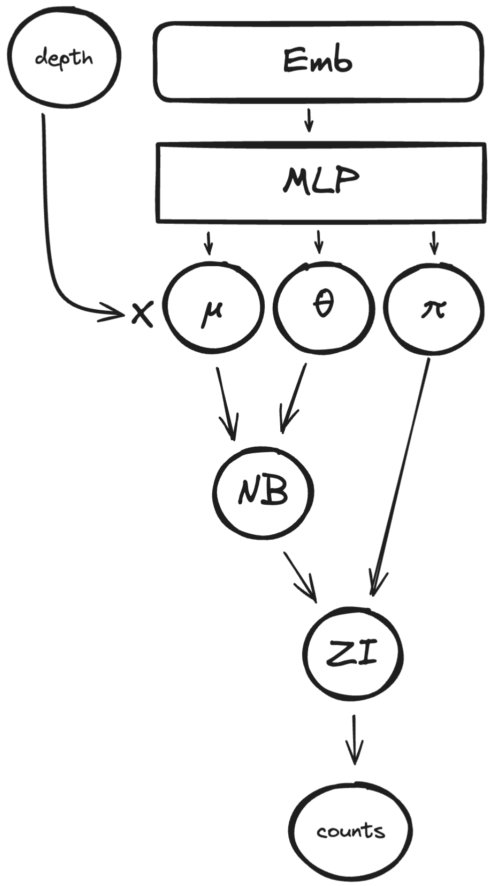

Schematic representation of the zero-inflated negative binomial graphical model of the expression decoder. We generate three values $\mu$, $\theta$, $\pi$ which are used to model a distribution. We also multiply the $\mu$ with the depth (or total count) over the cell.

#### FIG S13: Hierarchical classifier

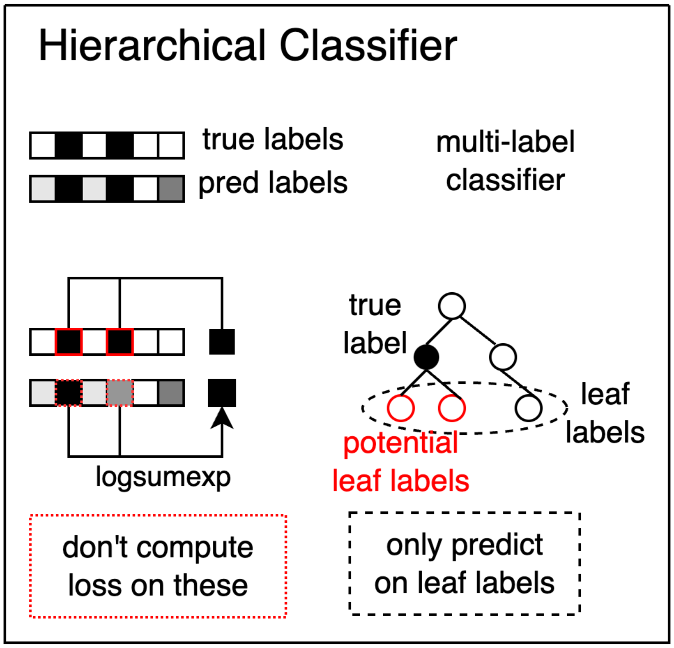

Schematic representation of the hierarchical classifier and its behavior during training. We can train on labels not predicted by the classifier as long as they are parent to one of the predicted labels in the ontological tree.

#### FIG S14: Detailed representation of the bottleneck learning procedure

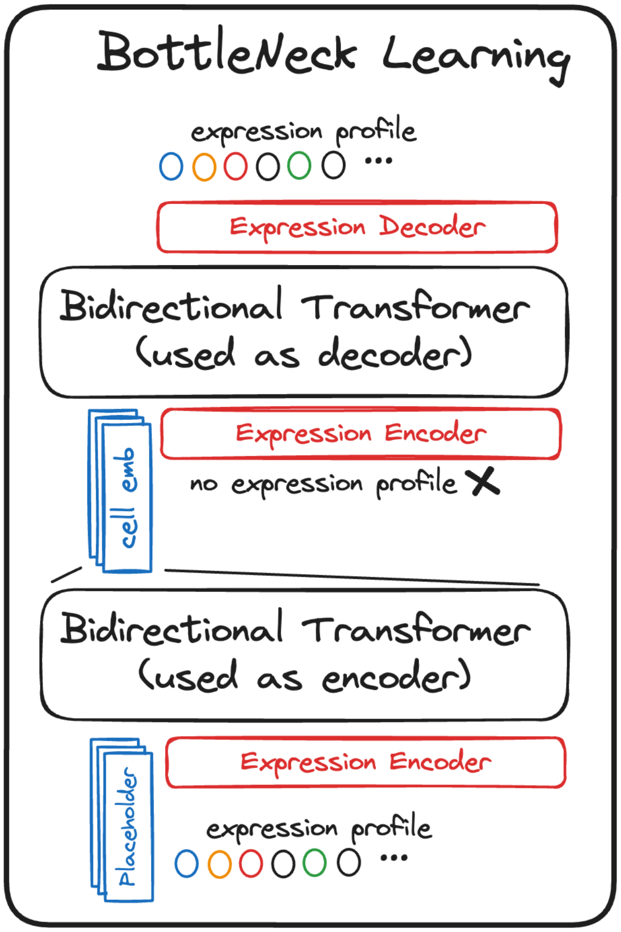

Schematic representation of the bottleneck learning procedure where scPRINT's Bidirectional Transformer Encoder is used both as the "Encoder" and "Decoder" of an auto-encoding (AE) bottleneck learning scheme.

### FIG S15: Schematic representation of our dataloader

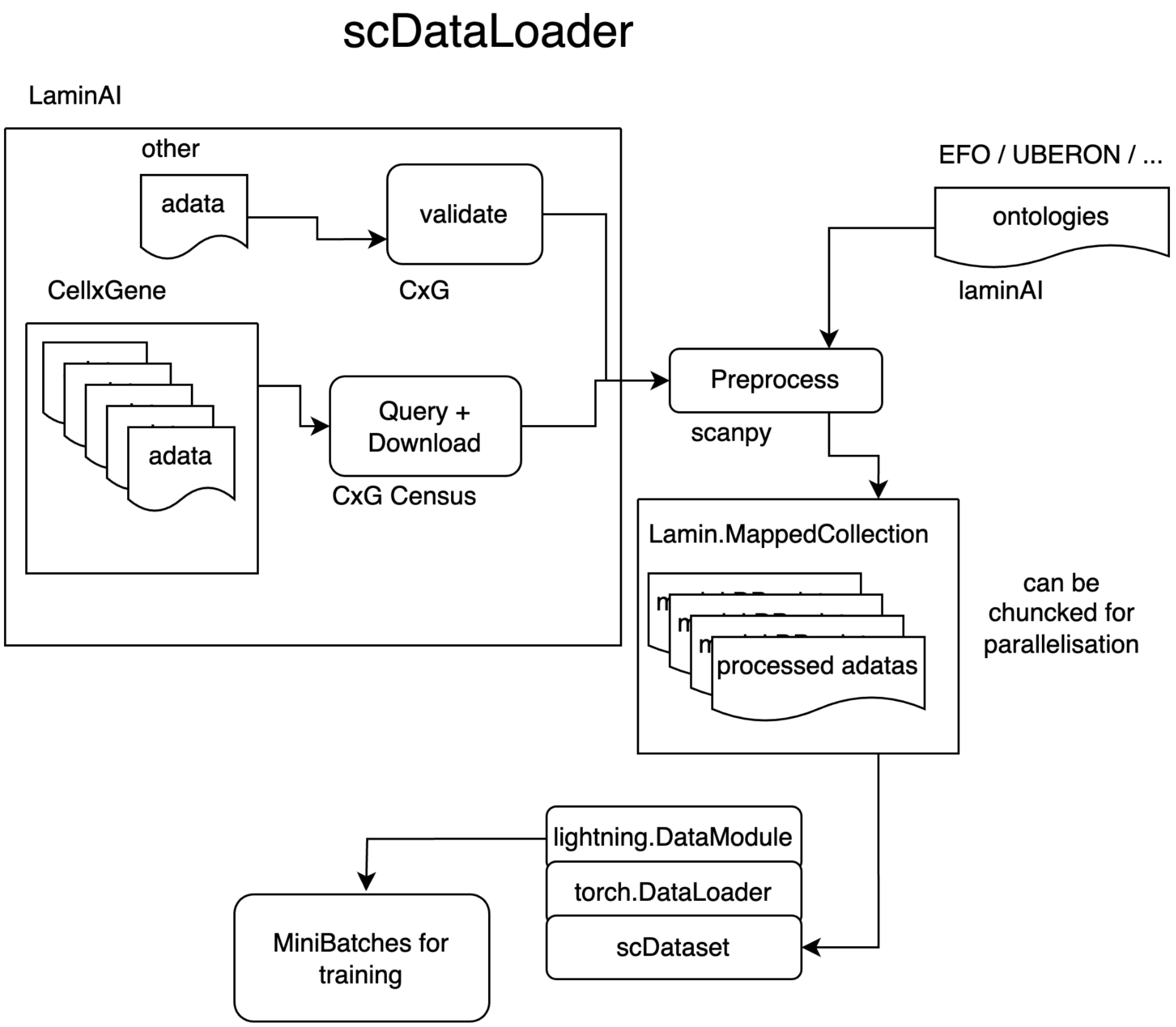

Schematic representation of scDataLoader. Using Lamin.ai, we download and preprocess all cellxgene datasets as AnnDatas. We can also add and validate other expression datasets using lamin.ai. Based on lightning's datamodule framework, torch's dataloaders, our weighted random sampler, and lamin.ai's mapped collection, we can then sample minibatches for pre-training across thousands of datasets and millions of cells with weighted random sampling.
